# Temporally resolved growth patterns reveal novel information about the polygenic nature of complex quantitative traits

**DOI:** 10.1101/2024.06.29.601327

**Authors:** Dorothy D. Sweet, Sara B. Tirado, Julian Cooper, Nathan M. Springer, Cory D. Hirsch, Candice N. Hirsch

## Abstract

Plant height can be an indicator of plant health across environments and used to identify superior genotypes or evaluate abiotic stress factors. Typically plant height is measured at a single time point when plants have reached terminal height for the season. Evaluating plant height using unoccupied aerial vehicles (UAVs) is faster, allowing for measurements throughout the growing season, which facilitates a better understanding of plant-environment interactions and the genetic basis of this complex trait. To assess variation throughout development, plant height data was collected weekly for a panel of ∼500 diverse maize inbred lines over four growing seasons. The variation in plant height throughout the season was significantly explained by genotype, year, and genotype-by-year interactions to varying extents throughout development. Genome-wide association studies revealed significant SNPs associated with plant height and growth rate at different parts of the growing season specific to certain phases of vegetative growth that would not be identified by terminal height associations alone. When plant height growth rates were compared to growth rates estimated from canopy cover, greater Fréchet distance stability was observed in plant height growth curves than for canopy cover. This indicated canopy cover may be more useful for understanding environmental modulation of overall plant growth and plant height better for understanding genotypic modulation of overall plant growth. This study demonstrated that substantial information can be gained from high temporal resolution data to understand how plants differentially interact with the environment and can enhance our understanding of the genetic basis of complex polygenic traits.

## Introduction

Advancements in crop improvement and production are essential to keep up with the current rates of population growth and changing climate. Over the past few decades, progress in sequencing technology has driven the development and implementation of genomic sequencing and marker platforms that can speed up crop improvement (Yang *et al*., 2020). Advances in the collection of accurate, high resolution phenotypic traits across many crop varieties and environments are also essential to meet this demand. Most current methods of trait acquisition are time-consuming, laborious, and only collected at a single time point at the end of the season, which leaves high-throughput phenotyping in field environments as a major bottleneck in crop improvement (White *et al*., 2012; Silva-Perez *et al*., 2018).

Plant height is an important trait in maize (*Zea mays* L.) breeding programs and can be used to monitor growth rates (Wang *et al*., 2018), assess plant health (Dhami, 2019), and predict yield (Boomsma *et al*., 2010; Yin, McClure, *et al*., 2011). Measurements of plant height have also been used to determine field spatial variability when evaluating abiotic influences such as nitrogen content (Yin, Jaja, *et al*., 2011; Katsvairo *et al*., 2003). These findings suggested plant height and estimated growth rates can be used as metrics to identify agronomically superior cultivars in plant breeding programs and to develop management practices to account for spatial variation in production fields. Unoccupied aerial vehicles (UAVs) are particularly useful for collecting maize plant height information since manual collection using rulers is not only time-consuming and often only collected at the end of the growing season, but is also associated with error due to viewing angles of data collectors or random sampling error (Tirado *et al*., 2020). UAVs can also collect data when issues such as wet soil and plant lodging prevent ground vehicles from navigating plots (Tirado *et al*., 2020).

Detection of plant height from UAV imagery has advanced through the use of structure from motion (SfM) which can create 3D reconstructions from overlapping overhead images (James and Robson, 2012). The 3D reconstructions developed through SfM can be used to create orthomosaics and digital elevation models which can be used to extract traits such as plant height. Multiple examples of accurate plant height estimates using UAV imagery have been shown in crops such as barley (*Hordeum vulgare* L.) (Bendig *et al*., 2014; Herzig *et al*., 2021), cabbage (*Brassica oleracea* var. *capitata* L.) (Moeckel *et al*., 2018), cotton (*Gossypium hirsutum* L.) (Feng *et al*., 2019; Liu *et al*., 2020), eggplant (*Solanum melongena* L.) (Moeckel *et al*., 2018), faba bean (*Vicia faba* L.) (Ji *et al*., 2022), maize (Anthony *et al*., 2014; Geipel *et al*., 2014; Grenzdörffer, 2014; Su *et al*., 2019; Varela *et al*., 2017; Anderson *et al*., 2019; Malambo *et al*., 2018; Shi *et al*., 2016; Tirado *et al*., 2020; Adak, Murray, Božinović, *et al*., 2021; Letsoin *et al*., 2023), potato (*Solanum tuberosum* L.) (de Jesus Colwell *et al*., 2021; Xie *et al*., 2022; Njane *et al*., 2023), rapeseed (*Brassica napus* L.) (Xie *et al*., 2021), sorghum (*Sorghum bicolor* L.) (Chang *et al*., 2017; Shi *et al*., 2016; Watanabe *et al*., 2017; Gano *et al*., 2021), soybean (*Glycine max* L.) (Li *et al*., 2022), tomato (*Solanum lycopersicum* L.) (Moeckel *et al*., 2018), and wheat (*Triticum aestivum* L.) (Holman *et al*., 2016; Madec *et al*., 2017; Michalski *et al*., 2018; Volpato *et al*., 2021). Studies on plant height using UAVs have shown variable levels of success when compared to manual measurements due to plant structure, field layout, and improvements in best practices as more research was completed (Holman *et al*., 2016; Sweet *et al*., 2022).

A major advantage to using UAVs to measure traits such as plant height is the ability to collect trait data at regular intervals throughout the growing season. High temporal resolution data collection facilitates a better understanding of end of season traits that are the culmination of a growing season’s worth of genotype-by-environment interactions (de Jesus Colwell *et al*., 2021). Such temporal measurements have been completed on canopy cover and biomass in soybean (Moreira *et al*., 2019; Herrero-Huerta and Rainey, 2019; Li *et al*., 2022; Freitas Moreira *et al*., 2021), plant height, canopy cover, growth dynamics, and yield prediction in barley (Herzig *et al*., 2021), plant height and canopy cover in banana (*Musa paradisiaca* L.) (Aeberli *et al*., 2023), nutrient status, canopy cover, canopy volume, and plant height in potatoes (Liu *et al*., 2021; de Jesus Colwell *et al*., 2021), plant height in cotton (da Silva Andrea *et al*., 2023), and plant height in eggplant, tomato, cabbage (Moeckel *et al*., 2018). In maize, multi-temporal measurements of multiple traits have been used to explain the effect of phosphorus on plant growth and final yield (Pedersen *et al*., 2021; Herrmann *et al*., 2020), phenotypic variation due to genotypic background differences (Pugh *et al*., 2018; Han *et al*., 2019; Alper Adak, Anderson, *et al*., 2023; A. Adak *et al*., 2023; Adak, Murray, Božinović, *et al*., 2021; Adak *et al*., 2024), phenotypic variation due to environmental factors such as drought (Machado *et al*., 2002) or high-speed wind events (Tirado *et al*., 2021), and canopy cover (Jin *et al*., 2020; Lu *et al*., 2021).

In addition to capturing phenotypic responses to the environment, high temporal resolution phenotyping can facilitate a more complete understanding of the genetic factors controlling a trait. Previous research looking at the genetic factors controlling maize plant height was completed with single measurements of terminal plant height due to the labor intensity of manual height collection (Peiffer *et al*., 2014). A study evaluating terminal plant height of many maize inbred lines in multiple environments found plant height was highly heritable and predictable with models, but also highly polygenic with many small effect alleles contributing to the overall height which are difficult to identify (Peiffer *et al*., 2014). Subsequent studies with higher temporal data collection were able to identify QTLs associated with early and mid season plant height, growth rates, and growth curves (A. Adak *et al*., 2023; Wang *et al*., 2019; Adak, Conrad, *et al*., 2021). These studies varied in the number of timepoints at which plant height was measured, the genetic diversity of the material used, and the number of environments evaluated. For example, Adak et al. (2023) collected RGB images of 280 maize hybrids over 15 timepoints, and identified 241 genome-wide association study (GWAS) peaks over 36 temporal phenotypes (A. Adak *et al*., 2023). These early studies provided valuable information in understanding of genetic variation in growth rates, yet understanding of the impact of genotype, environment, and genotype-by-environment interaction (GxE) on temporally resolved growth curves is still lacking due to the limited number of growth environments and/or genotypes included in the studies.

Here we report on high temporal resolution phenotyping of a panel of over 500 diverse maize lines from the Wisconsin diversity panel (Hansey *et al*., 2011) that represent the breadth of variation in temperate maize germplasm grown over four growing environments between 2018-2021. The objectives of this study were 1) to determine how much plant height variance is explained by different experimental factors (i.e. genotype, environment, and GxE) throughout development, 2) to identify patterns of growth rates within and between growing seasons and assess the consistency of these growth patterns, 3) to map the genetic basis of plant height and growth rate throughout vegetative growth across multiple environments, and 4) to evaluate the differences and relative utility of using different traits (i.e. plant height versus canopy cover) to generate plant growth curves.

## Results and Discussion

### LOESS curves fit to the data break the growing season into 3 phases

Flights were conducted at 14 timepoints in 2018, 27 timepoints in 2019, 12 timepoints in 2020, and 11 timepoints in 2021 to obtain plant height data throughout each growing season (Table S1 and Table S2). In order to directly and quantitatively compare growth curves among genotypes and years, a method of standardization was needed to align similar time points from one year to the next. Converting the dates of flights to growing degree days (GDDs) made it possible to compare growth stages across years while accounting for different planting dates and temperatures across the years (Table S3). LOESS curves were fit to the data from each plot, and plant height in 50 GDD increments throughout the growing season were predicted from the curves (Figure S1 and Table S4). This allowed for direct comparisons regardless of the actual flight dates across the years. The growth curves were annotated to denote lag phase, exponential growth phase, and terminal height (Figure 1a). This dissection of the growing season is important as both plant growth and grain yield are differentially affected by temperature and precipitation throughout the growing season, especially when the variation occurs early in the season (Dodig *et al*., 2021; Butts-Wilmsmeyer *et al*., 2019; Claassen and Shaw, 1970).

**Figure 1.**
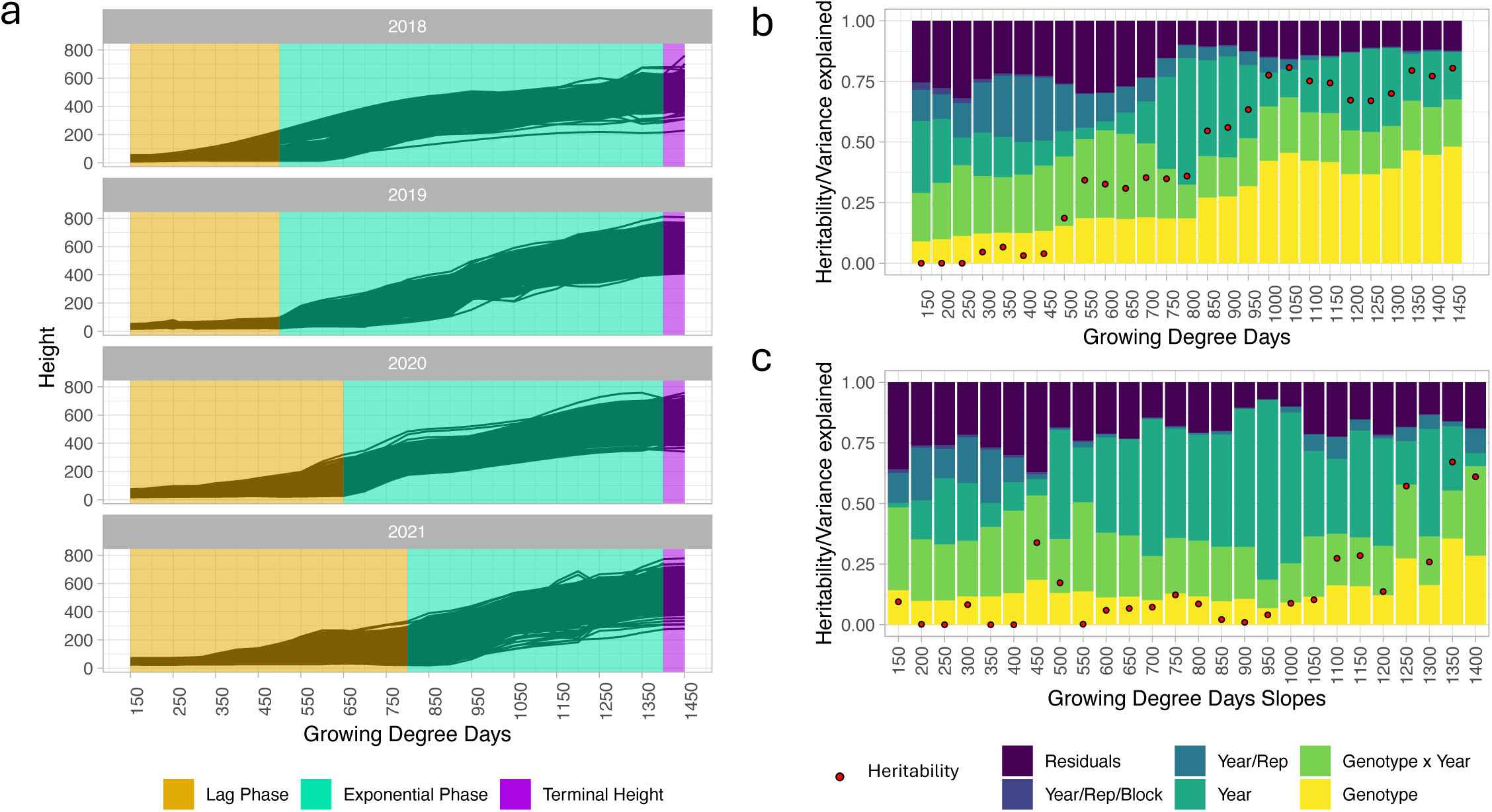
Temporal plant height and growth rate analysis of variance. (a) LOESS curves of plant height broken into three phases of the growing season based on the performance of genotypes in each year. (b) Variance explained and heritability at each plant height time point throughout the growing season. (c) Variance explained and heritability at each growth rate time point throughout the growing season.

The beginning of the exponential growth phase differed between years, with 2018 and 2019 reaching the phase as early as 500 GDDs and 2021 reaching exponential growth as late as 800 GDDs (Figure 1a). This is likely due to differences in the speed of GDD accumulation, early season precipitation, and planting time. The planting dates in 2018 and 2019 were later (05/14/2018 and 05/30/2019), while 2020 and 2021 were planted earlier in the calendar year (05/07/2020 and 05/06/2021) allowing for a more gradual accumulation of GDDs (Figure S1b and Table S5). The most precipitation prior to the exponential growth period was accumulated in 2020 with 6.17 inches while the rest of the years had accumulated only 3.05, 3.17, and 3.56 inches in 2018, 2019, and 2021, respectively (Table S5). Additionally, 2021 had a long dry spell (22 days with minimal to no rain) from 219 to 724 GDDs while other years showed no significant water stress during that time.

### Heritability of plant height increased over time, while growth rate was largely explained by year for much of the growing season

Highly variable plant height and growth rate values across years indicated that aspects of the environment had a large phenotypic impact in addition to the genetic background of the material. An analysis of variance (ANOVA) showed that the percent variance explained by genotype gradually increased throughout the growing season, which coincided with increased heritability throughout the season (Figure 1b). Work using temporal plant height in a panel composed of commercial and experimental hybrid and inbred maize genotypes completed by Pugh et al. (2018) showed similar trends in percent variance explained by genotype. In contrast, replicate explained the greatest proportion of the observed variation early in the growing season starting at 200 GDD due to highly variable soil temperature and moisture conditions early in the growing season resulting in more spatial variation within the field (Figure 1b). As the growing seasons continued there was almost no variation explained by replicate, and by 1150 GDDs the percent variation explained (PVE) for replicate did not exceed 0.8% for the remainder of the growing season. Pugh et al. (2018) also showed large variation due to replicate earlier in the season for maize planted at the optimal time that decreased as the season continued. Genotype-by-year interaction also explained more variance early in the growing season when the difference in days to accumulate early GDDs was highly variable (Figure S1b). Throughout the exponential growth phase, a larger portion of the variance partitioned to the year term than any other time in the growing season. During this phase of vegetative growth, stalk elongation can be severely affected by environmental factors such as water availability (Claassen and Shaw, 1970). Indeed, the amount of water accumulation through this phase of the growing season differed nearly 3 fold over the four seasons this experiment was grown, ranging from 2.47 inches in 2021 to 6.78 inches in 2018 (Minnesota Department of Natural Resources, n.d.) (Table S5).

Rate of growth, calculated as the slope between plant heights, generally had less distinctive patterns than plant height in how variance partitioned (Figure 1c). Overall, a much larger proportion of the variation was explained by year and genotype-by-year interaction, demonstrating that growth rate was less dependent on the genotype and influenced more by the environment. This was particularly evident during the exponential growth phase when 17-74% percent of the total variation was explained by year. Similar to plant height, the highest PVE by replicate for growth rate also occurred early in the season when soil temperature and other soil conditions were more variable.

### Clustering revealed three growth patterns that fluctuated within genotype from year to year

The ANOVA partitioned variation at each individual time point. To further understand the significant variance explained by year and genotype-by-year interaction on the full growth curve, the growth curves were clustered within each year using fuzzy c-means clustering and patterns were compared within and among years. Fuzzy c-means clustering places each curve into each cluster with various degrees of fit into the clusters ranging from 0-1 (Table S6). This approach allowed curves to be grouped into defined clusters based on wellness of fit, while also providing information on how similar the genotypes within a group are to each other (Han *et al*., 2019). The number of clusters was chosen based on the distribution of the largest wellness of fit value for each curve (Figure S2) with 3 clusters having a fairly normal distribution. The three clusters were characterized as genotypes that start tall relative to other genotypes in the experiment and end tall, genotypes that start short and end short, and genotypes that start short and end tall (Figure 2A), with some years fitting these patterns better than others (Figure S3). In order to compare groups across years, each curve was assigned to a group based on its largest wellness of fit value. A comparison of distinct curves across all four years showed that genotypes do not always have the same growth pattern (Figure 2b, Table S7). The short to short pattern kept the most consistent genotypes with 39 that are present in all four years out of the 403 genotypes with that pattern in at least one year. Despite the low percentage of genotypes with the same pattern across all four years, the high recurrence of genotypes within single pairings showed that some years are more similar than others. For instance, between 2018 and 2020, 111 genotypes stayed in the short to short growth pattern out of the 158 genotypes in that pattern in 2018, meaning 70% of the genotypes with a short to short pattern in 2018 had a short to short pattern in 2020 (Figure 2b).

**Figure 2.**
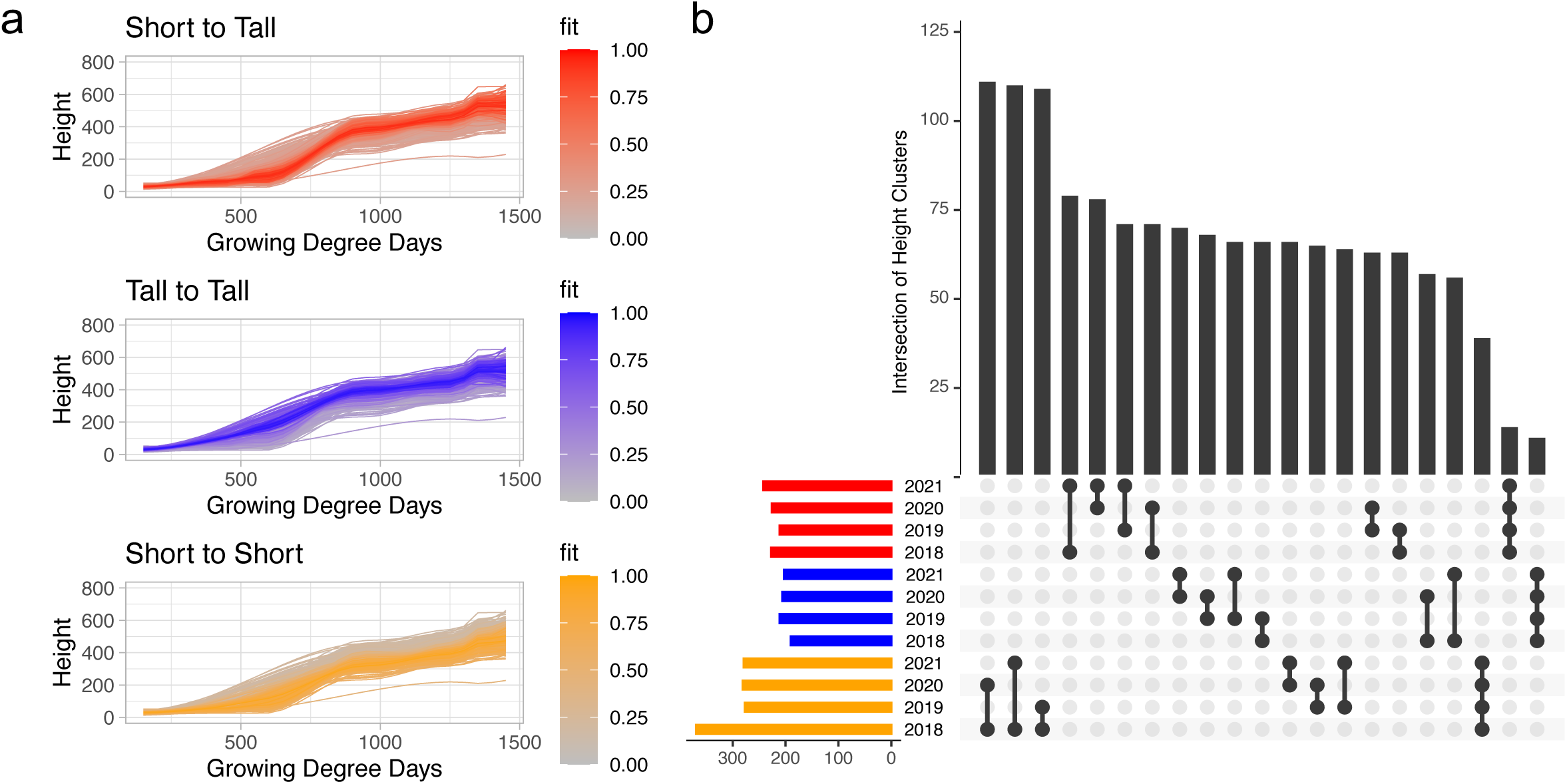
Fuzzy c-means clustering of LOESS growth curves. (a) Fuzzy c-means clusters of 2018 LOESS growth curves with shading equating wellness of fit for each curve into the specified cluster. (b) Upset plot showing overlap of genotypes between growth curve clusters across years. Each curve is classified into a cluster based on the highest fit for each curve. Cluster types are represented by identifying colors (yellow: tall to tall; blue: short to short; red: short to tall).

### Fréchet distances of growth curves provided more information on plasticity and stability than terminal height

Clustering the genotypes into groups gave an idea of how similar the growth curves were between different genotypes, but we were also interested in a quantitative measurement of the similarity of the curve for each genotype across years. Other studies have employed practices such as ‘phenotypic similarity trees’ to calculate similarity distances based on many independent phenotypic traits (Chen *et al*., 2014). As our data is a continuous series of data points that make up a growth curve we instead evaluated the curve similarity using the Fréchet distance. The Fréchet distance is a measurement of similarity that takes location and order of points along the curve into account and is often described as the shortest leash possible when a person walks on one curve and their dog walks on the other (Fréchet, 1906). In this case, it can be used to observe general trends between years, as well as the plasticity of specific genotypes across the years. To assess overall differences among years, each year was compared to every other year with average Fréchet distances over all genotypes varying remarkably (Figure 3a, Figure S4, and Table S8). Eighteen environmental parameters in daily increments throughout the growing season were used to calculate pairwise environmental correlations between each combination of years (Table S5). A strong relationship between the pair-wise average Fréchet distances and the pair-wise environmental correlations was observed (Pearson correlation coefficient = 0.80; Figure S4)). The very large average Fréchet distance between 2018 and 2019 was likely due to these years having the largest difference in GDD accumulation across all four years. In 2018, GDDs began to accumulate earlier, and by mid to late August there is a large difference in the number of GDDs accumulated (August 17, 2018 - 2248.5 GDDs, August 21, 2019 - 1650.5 GDDs) (Figure S1b). Conversely, in 2020 and 2021, which has the smallest average Fréchet distance across year pairs (0.3650), there is very similar accumulation of GDDs relative to calendar day in the growing season (July 22, 2020 - 1340 GDDs, July 23, 2021 - 1421 GDDs). This indicates that while GDDs are a good indicator of developmental stages, they are not necessarily linked to plant growth.

**Figure 3.**
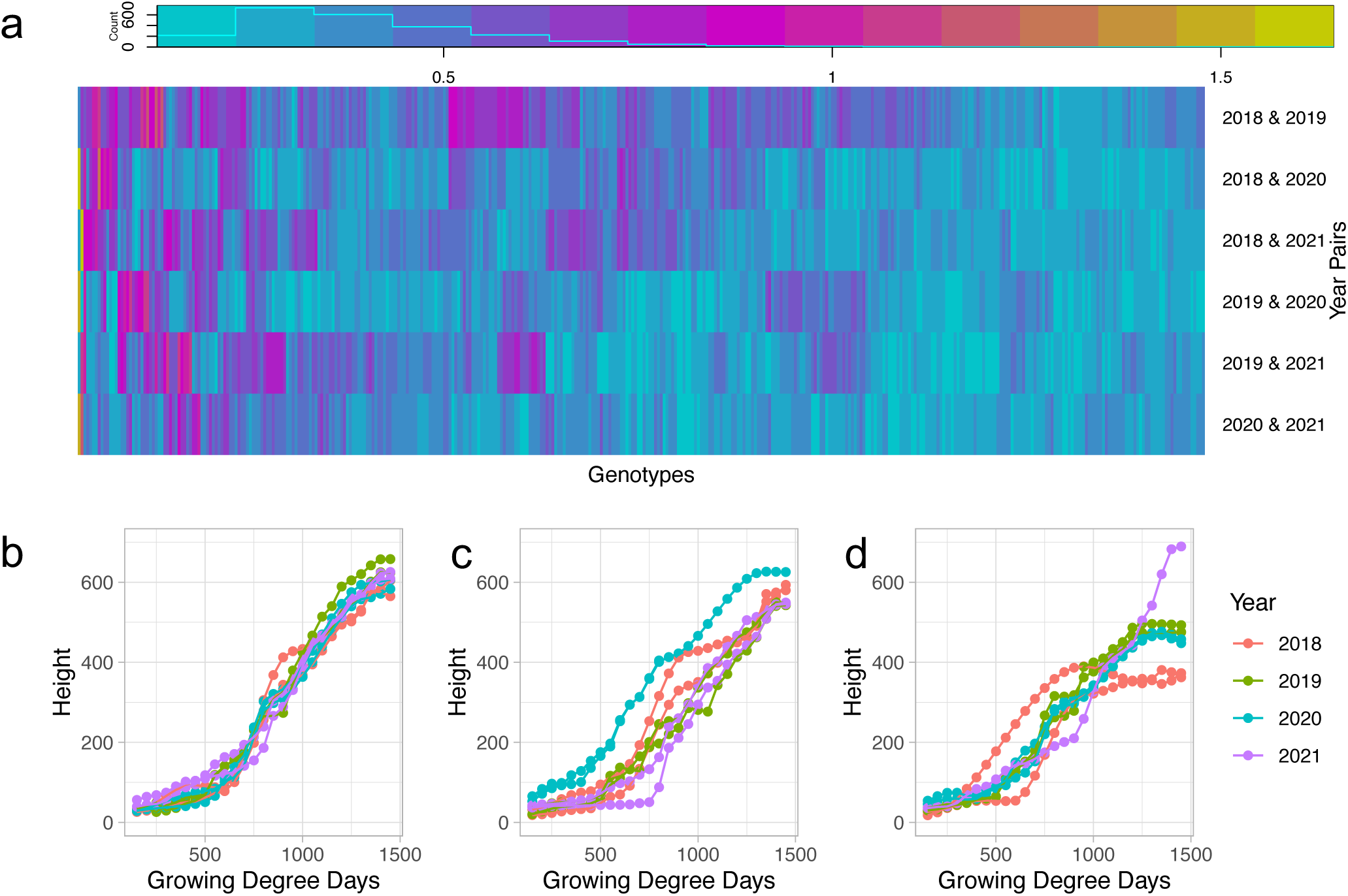
Fréchet distances for each genotype present in all four years. (a) Pairwise Fréchet distance values comparing plant height growth curves of the same genotype across different years. (b) Example genotype (PHG47) representing genotypes with consistently low Fréchet distance values and low variation in terminal height. (c) Example genotype (YING-55) representing genotypes with high Fréchet distance values and low variation in terminal height. (d) Example genotype (PHN66) representing genotypes with high Fréchet distance values and high variation in terminal height.

Fréchet distances were also used to assess differences in overall growth curve topology within genotypes across year pairs as a metric of growth stability for individual genotypes. A wide range of Fréchet distances were observed across all of the genotypes (average 0.209-1.055 across all year pairs, Table S8), reflecting variable environmental responsiveness of the different genotypes in this population throughout vegetative development. To determine the impact of variable growth patterns across years on terminal height the Fréchet distances across years were compared to variance in terminal plant height. The average variance of terminal plant height across years ranged from 4 to 13035, also reflective of the variability in environmental responsiveness across the genotypes. While a wide range of variance in both growth curve and terminal height stability was observed, there was not a strong relationship between average stability across environments for growth curve and terminal plant height (Pearson correlation coefficient = 0.36). Thus the relative variance in end of season traits does not always reflect the variance in the growth and development required to reach the end of season trait. Some genotypes were stable for both growth curves (low Fréchet distance) and variance in terminal plant height (e.g. Figure 3b). Of the 100 most stable genotypes for terminal plant height (20th percentile), 26 of these were also in the 20th percentile for growth curve stability. In contrast, other genotypes had very stable terminal height, but their growth curves varied substantially between growing seasons (e.g. Figure 3c). Of the same 100 most stable genotypes for terminal plant height, 10 of these were in the top 80th percentile for Fréchet distance, and did not reach the consistent terminal height in the same way each year. Most (n=39) of the remaining genotypes with highly unstable growth curves (top 80th percentile) resulted in high variability in terminal height (top 80th percentile) (e.g. Figure 3d), and would likely be identified as having high plasticity across these growth environments regardless of analysis with terminal plant height or full growth curves. This analysis demonstrates that full growth curves give different, if not more complete, information about genotypic plasticity than terminal height, which has often been used (Mu *et al*., 2022).

### GWAS identified 669 significant SNPs throughout the growing season that were not identified at terminal height

To identify the genetic basis of the observed variation in growth curves across the genotypes in this study, genome-wide association studies (GWAS) were conducted on plant height and growth rate at 50 GDD windows throughout the growing season. Other studies have completed temporal GWAS with plant height in maize (Wang *et al*., 2023; Adak, Conrad, *et al*., 2021; Farfan *et al*., 2015) as well as vegetative indices (Adak, Murray, Anderson, *et al*., 2021; Alper Adak, Kang, *et al*., 2023; Wang *et al*., 2021; Rodene *et al*., 2022), however, this study is unique by using a large diverse inbred panel in field settings over multiple years. This experimental design allowed a deeper understanding of how the genetics of complex traits such as plant height interact with the environment. GWAS completed for each plant height GDD, growth rate, and extracted terminal plant height revealed a total of 717 non-redundant unlinked SNPs (Table S9). Of these SNPs, 48 were identified at terminal height (Figure 4a), with the majority of these 48 (n=42) not identified at any other time point. Thus, the vast majority of the significant SNPs identified in this study (n=669), would be missed if the GWAS was conducted only at the terminal time point as has been previously done (Zhang *et al*., 2019; Wallace *et al*., 2016; Weng *et al*., 2011). SNPs that were not identified in terminal height, but were identified earlier in the season may be helpful in improving early season plant resilience to detrimental weather effects such as low temperatures or high rainfall and may allow plants to outgrow competing weeds. Significant SNPs identified in early growth rates (Figure 4b) may be especially important as these SNPs appear to be particularly affected by the environment.

**Figure 4.**
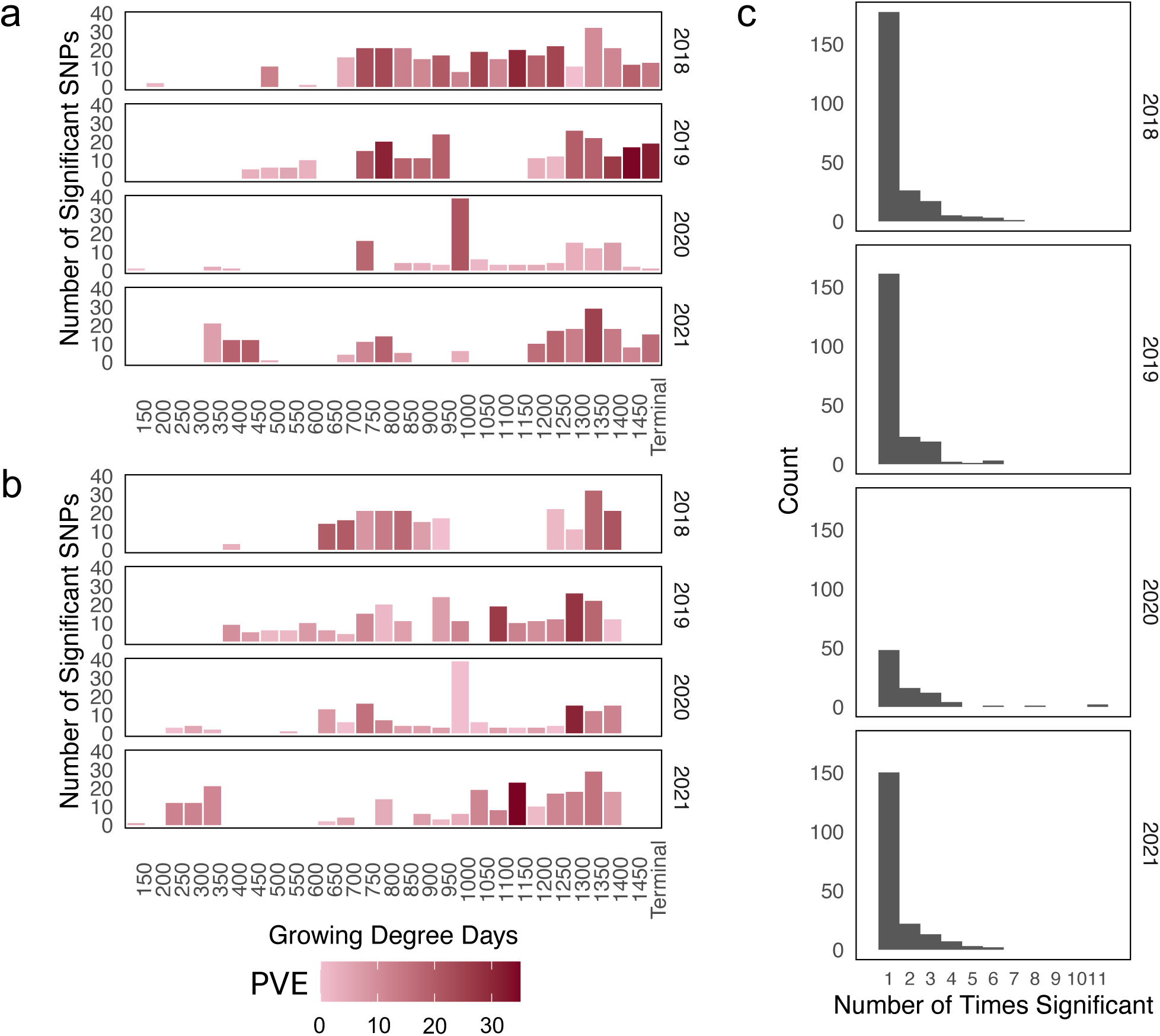
Significant SNPs identified with GWAS across development. (a) Results from plant height GWAS at each growing degree day interval throughout the growing season shaded by percent variance explained (PVE) of the significant SNPs at that time point. (b) Results from growth rate GWAS at each growing degree day interval throughout the growing season shaded by PVE of the significant SNPs at that interval. (c) Number of time points (height) or intervals (rate) in which a SNP was identified as significant within a year.

Plant height has long been documented as a complex, quantitative trait, even in studies that only assessed terminal variation (Zhang *et al*., 2019; Wallace *et al*., 2016; Weng *et al*., 2011). The large number (n=717) of significant SNPs throughout the growing season in this study demonstrated further complexity in this trait than previously recognized with other studies identifying between 4 and 204 loci associated with plant height (Mazaheri *et al*., 2019; Peiffer *et al*., 2014; Zhang *et al*., 2019; Weng *et al*., 2011; Adak, Murray, Anderson, *et al*., 2021; Wang *et al*., 2019). Of the 717 significant SNPs identified in this study, 22 overlapped with previously identified plant height QTL (Table S10) (Adak, Murray, Anderson, *et al*., 2021; Zhang *et al*., 2019; Wang *et al*., 2019; Mazaheri *et al*., 2019). These overlapping QTL likely represent highly stable QTL with minimal QTL-by-background or QTL-by-environment effects. The limited number of loci in previous studies that did not overlap the loci identified in the current study could be false positives in those studies, loci that are not segregating in the germplasm used in the current study and so unable to be detected, or have a strong QTL-by-background or QTL-by-environment interaction. Conversely, there were substantially more novel loci identified in our study compared to these previous studies that were only able to be identified due to the use of high resolution temporal data.

To assess how much of the total variation was captured by the SNPs that were identified as significant, the total PVE by the subset of significant SNPs at each time point was calculated. The PVE at any given time point-by-year combination ranged from 1.5% to 34.6% for height (Figure 4a) and from 0.0% to 27.8% for growth rate when looking at SNPs significant only at that individual time point-by-year combination (Figure 4b). PVE by significant SNPs at a single time point tended to explain more variance mid to late season in plant height (Figure 4a), which is consistent with the observed increase in heritability throughout the growing season (Figure 1b). We were also interested in the PVE at each time point using all significant SNPs across development to determine if these additional markers were able to explain substantially more variation. Indeed, PVE at any given time point-by-year combination calculated using all significant SNPs identified across time was much larger with PVE for height ranging from 2.0% to 70% and from 0.6% to 60.6% for growth rate. This demonstrates that high temporal resolution phenotyping facilitates a deeper understanding of the complete genetic basis of this complex trait.

We were interested to know how many of the identified SNPs were closely tied to environmental interactions and if SNPs were impacting plant height for long periods of time. In order to ascertain this information, we looked into the number of times SNPs were found to be significant within years and if the same SNPs were significant across years. Of the total 717 significant SNPs, 175 were found to be significant at multiple time points, leaving most of the significant SNPs to only appear once (Figure 4c, Table S10). The 175 SNPs appeared from 2 to 11 separate times with most reappearances occurring within the same year, including 2 SNPs in 2020 that were significant at 11 different time points throughout development. Seven of these SNPs found at multiple time points in a single year overlapped with previously identified regions (Mazaheri *et al*., 2019; Adak, Murray, Anderson, *et al*., 2021; Wang *et al*., 2019; Zhang *et al*., 2019), further demonstrating the stability of these loci not only across backgrounds and environments, but also throughout development. Six of the recurring SNPs were found to be significant over multiple years, five between 2018 and 2019 and one between 2019 and 2021. These SNPs may be less dependent on environmental factors and more generally impactful on growth. Two of these SNPs were found to be significant in multiple years toward the middle of the season during the exponential growth phase and are likely linked to stem elongation while four of them are significant at the end of the season and would be more likely to be tied to flowering time and picked up by studies only examining terminal height. Indeed, three of these four SNPs were also associated with terminal plant height.

The lack of overlap in significant SNPs over time and years is likely because the threshold for significance in GWAS studies is high (Fadista *et al*., 2016) and the effects of individual loci for our highly quantitative traits were relatively low (Figure 4a and b). Still, we were curious if patterns of effect sizes could be observed over time regardless of the significance of the marker at each time point as that may indicate importance at more time points regardless of statistical significance. For this analysis, all SNPs that were observed to be significant in at least one time point-by-year combination for either height (Figure 5a) or rate (Figure 5b) were assessed across all time points and years. Plant height (Figure 5a) showed a group of significant SNPs with gradually increasing effect size over the growing season (e.g. Figure 5a - Bracket 1, 5c), which may indicate these SNPs are important throughout growth, and contribute to overall terminal height. Some of these SNPs had the same effect size patterns over multiple years. Other SNPs had a larger effect size during mid vegetative growth without any large increases across the time point (Figure 5a - Bracket 2). As would be expected due to the few SNPs significant over multiple years, there are also a lot of SNP-year combinations that do not appear to have much of an effect at any point in the growing season.

**Figure 5.**
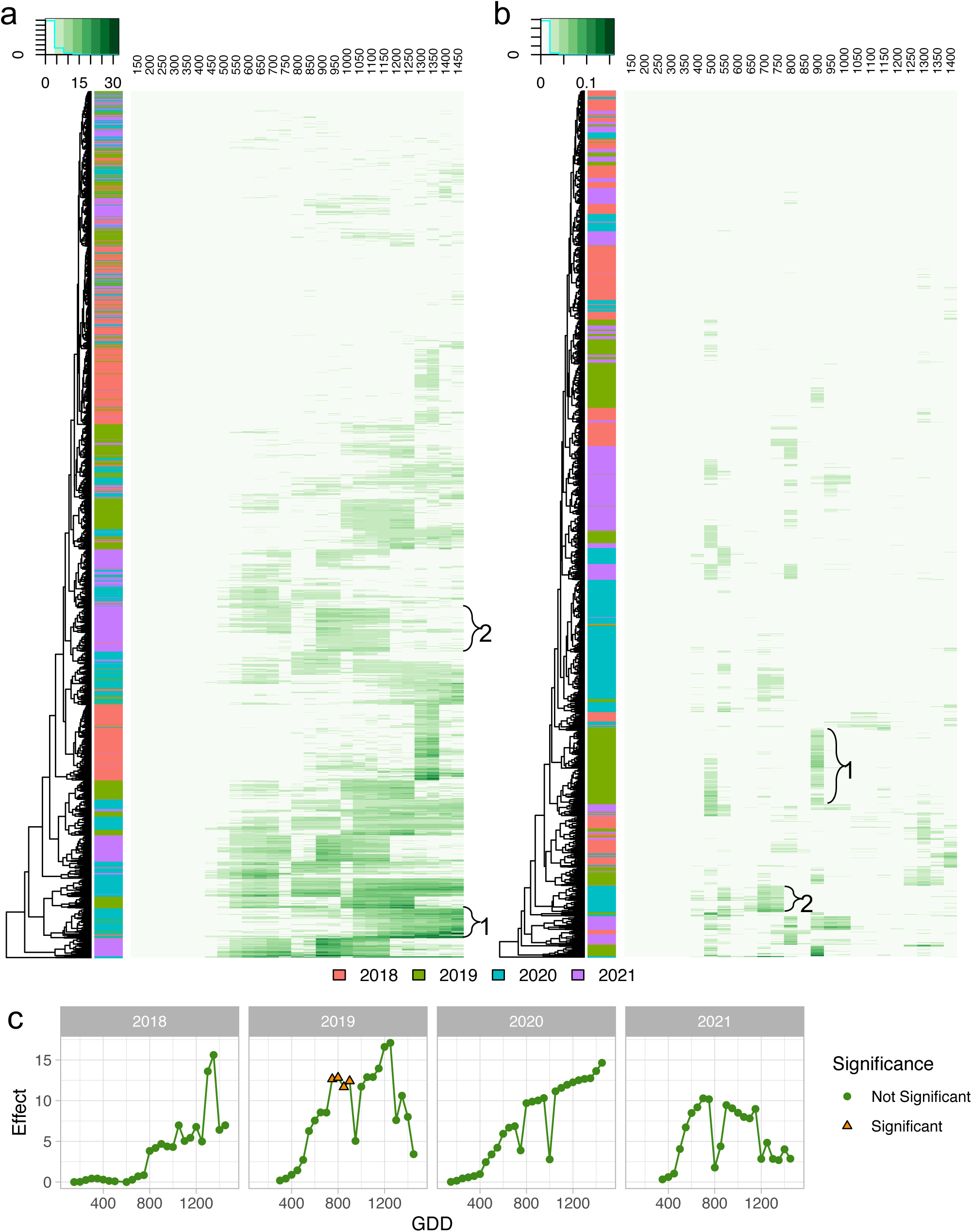
Significant SNP effects from GWAS conducted on plant height or growth rate across the growing season. (a) Absolute value of effect of significant SNPs from at least one time point-by-year assessed across all plant height GWAS iterations in every time point throughout the growing season in all four growing seasons. (b) Absolute value of effect of significant SNPs from at least one time point-by-year assessed across all growth rate GWAS iterations in every time point throughout the growing season in all four growing seasons. (c) Absolute value of effect of an example significant SNP from plant height located on Chr2 at position 169140523 in the B73 v4 reference genome assembly throughout each of the growing season.

The effect sizes of SNPs significantly associated with growth rate throughout the growing season had very different patterns from what was observed for height (Figure 5a and b). Growth rates clustered more by year and the effects were observed for very short periods of time without any indication of increasing or decreasing effect size throughout development. For instance, a group of SNP-year combinations from 2019 show their largest effect size at the 900-950 growth rate (Figure 5b - Bracket 1). This coincides with a lodging event that took place that year (Tirado *et al*., 2021). Another group of SNP-year combinations mostly from 2018 and 2020 had their largest effect from 700-800 GDDs which coincided with the early to mid-exponential growth phase in those years (Figure 5b - Bracket 2). These results are consistent with the ANOVA results in which a larger percentage of the observed variation was explained by year for growth rate (Figure 1c). The high variability of effect size linked to growth rate and the short time frames in which SNPs are significant highlight the extreme complexity of this trait and the major impact of immediate environmental conditions.

### Plant height is more genetically controlled, while canopy cover is more influenced by the environment

Canopy cover, or the percentage of ground covered by overhanging aboveground plant material, is another important trait contributing to yield potential, biomass accumulation, light interception, and weed suppression (Campillo *et al*., 2008; Jannink *et al*., 2001; Purcell, 2000; Xavier *et al*., 2017). In order to maximize the competitive benefits of canopy cover, most selections push toward higher cover percentages early in the growing season. Historical methods of measuring canopy cover are often time-consuming and difficult to complete (Purcell, 2000; Campillo *et al*., 2008). UAV collection of canopy cover increases both the speed and precision of collection by reducing the amount of time spent in the field and the objectivity of the observer and sun-angle (Purcell, 2000; Campillo *et al*., 2008). Similar to plant height, canopy cover increases throughout the vegetative growth stage, making it another potential trait for which growth patterns over developmental time can be assessed for genetic and environmental contributions.

To facilitate direct comparison of the growth patterns between plant height and canopy cover, comparable growth curves were generated for canopy cover data extracted from the same flight data that plant height was extracted (Figure S5)(Cooper *et al*., 2024), and values in 50 GDD windows throughout development were predicted from the curves (Table S11). Grouping the canopy cover growth curves using fuzzy c-means clustering revealed different types of patterns than were observed for plant height, with relative ranks among plots remaining more consistent throughout development (Figure S6). Differences in the Fréchet distances between the same genotypes over multiple years were also observed (Figure 3 and Figure S7), with larger average distances for canopy cover (average 0.575 for canopy cover versus 0.415 for plant height). This indicated that canopy cover is more influenced by interactions with environmental factors than plant height. Indeed, when variance was partitioned, the genotype-by-year interaction consistently explained 12-20% of the variation; while in plant height, genotype-by-year explained less than 10% of the variation for most of the growing season (Figure 1b and Figure S8). During the early exponential growth phase, year by itself explained the largest percentage of the variation in canopy cover while heritability was at its lowest indicating that environment regardless of genotype has a large impact on canopy coverage variation. Similarly to the Fréchet distance, the variation due to year suggested that canopy coverage is a good indication of the variation in environmental conditions present within the growing season.

### Plant height and canopy cover are not predictive of one another and contribute complementary information throughout the growing season

As canopy cover is more influenced by year and genotype-by-year interactions, it would be expected to observe differences in plant height and canopy cover progression throughout the season. To test this, direct comparisons of plant height and canopy cover were done. The correlation between plant height and canopy cover within a plot at each GDD interval was highly variable from year to year with a high correlation in early 2019 and 2021 and a low to negative correlation in early 2018 and mid 2020 (Figure 6a). These differences were likely due to environment variation that differentially affected plant height and canopy cover with the differences between plant height and canopy cover being genotype dependent and canopy cover overall being more variable. To further assess these differences, Fréchet distances were calculated between plant height and canopy cover within each plot (Figure 6b). While many genotype distances (n = 169) were consistent across years with a standard deviation less than 0.2, some genotypes showed higher values in some years compared to others, with 2019 having the largest distances. This result was unexpected as 2019 had a high to average correlation between plant height and canopy cover at individual points in time, but showed the largest individual Fréchet distances, which assess differences in the overall growth curve. This remained true when including all within plot comparisons regardless of presence in all four years. It is possible the Fréchet distance between plant height and canopy cover identified genotypes more affected by the lodging event that occurred during that year, or were more influenced by genotype-by-year interactions.

**Figure 6.**
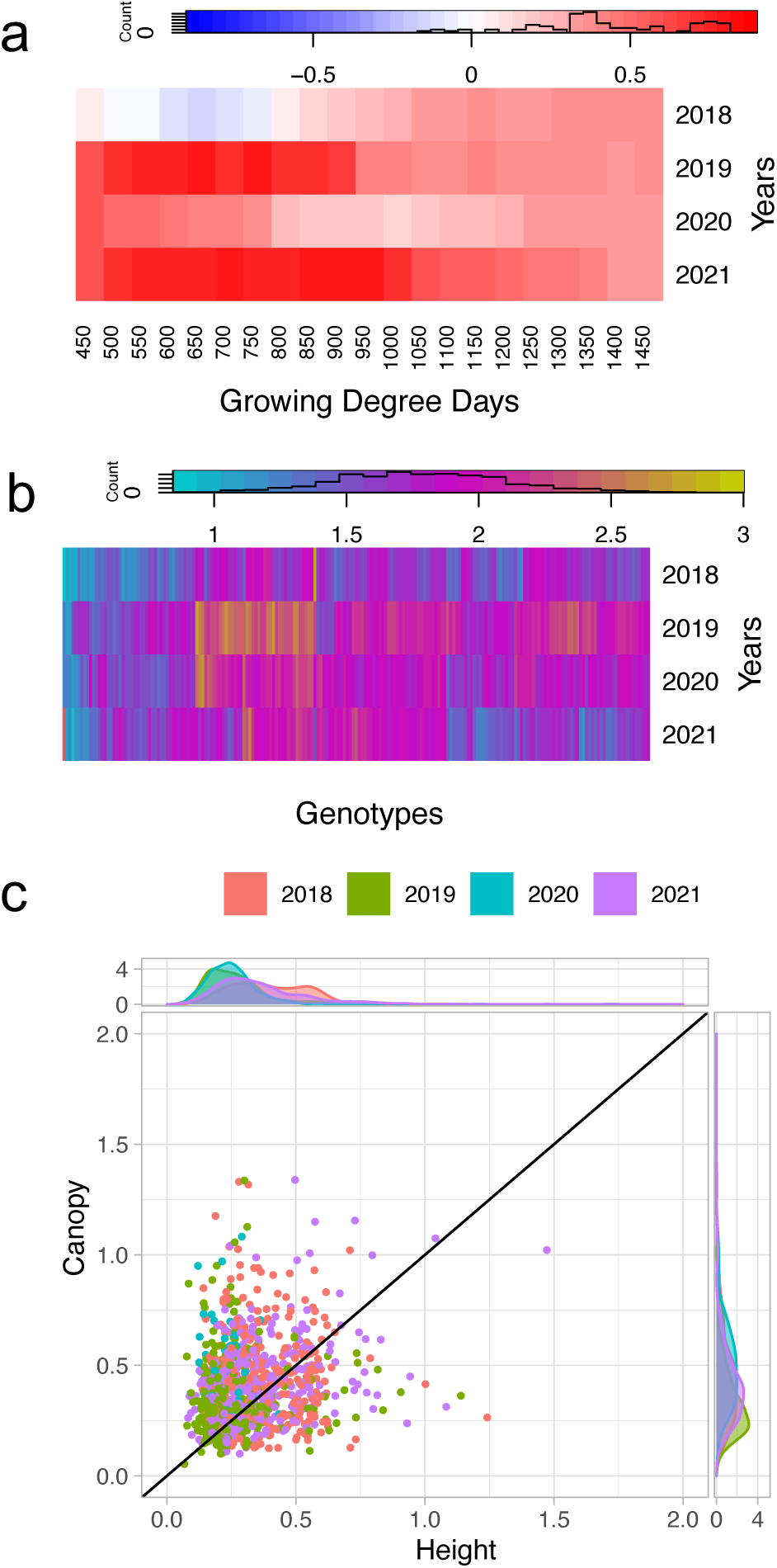
Relationships between plant height and canopy cover. (a) Pearson correlation between plant height and canopy cover across all plots at each GDD interval for each year. (b) Fréchet distance between the plant height growth curve and canopy cover growth curve within the same plot. (c) Fréchet distance between experimental entry replicates for plant height versus Fréchet distance between experimental entry replicates for canopy cover.

To further assess the difference in environmental responsiveness to within year spatial variation, we calculated Fréchet distances between replicates within a year for both plant height and canopy cover. Consistent with the between year observations for the same genotype in the same year, replicates had more similar growth curves for plant height than for canopy cover (Figure 6c). This supports that depending on the trait in which growth curves are made, different aspects of plant stability/plasticity will be captured, and that in this case canopy cover may be more beneficial in assessing plant responsiveness to environmental variation.

## Conclusions

Understanding both the genetic basis and phenotypic plasticity of complex traits such as plant height will only increase in importance as our climate becomes increasingly unpredictable. The advent of high-throughput phenotyping has and will continue to make the necessary data collection for such a task possible. In this study, we provided an in-depth analysis of the genetic and environmental factors that influenced plant height and growth rate in maize, as well as the methodology for understanding the genetic basis of temporal trait variation. The use of temporal traits in combination with metrics of stability, such as Fréchet distances, gave more information about genotypic plasticity than terminal traits alone, and was beneficial to understanding how plants respond to different environments. The genetic architecture of plant height was further underscored by numerous significant SNPs associated with plant height and growth rate throughout the growing season and illustrated the need for a better understanding of temporal traits and implementation for crop improvement.

Overall this study enhanced our understanding of the genetic basis of trait variation in maize by demonstrating the significant interplay between genetics and environment. The findings emphasize the importance of considering temporal dynamics in plant growth studies, which could inform breeding programs aimed at improving crop resilience and performance under varying environmental conditions.

## Materials and Methods

### Experimental Field Design

A set of 501 diverse inbred lines from the Wisconsin Diversity Panel (Hansey *et al*., 2011; Renk *et al*., 2021; Burns *et al*., 2021) were grown in the summers of 2018, 2019, 2020, and 2021. These trials were planted on May 14, 2018, May 30, 2019, May 7, 2020, and May 6, 2021 at the Minnesota Agricultural Experiment Station in St. Paul, MN. The lines were grown in single-row plots that were 19.5 feet long center-to-center, with 4 foot alleys, 30 inch row-spacing, and were planted at a density of approximately 70,000 plants per hectare. All experiments were planted as a randomized complete-block design with two replicates. Within replicates, genotypes were blocked by flowering time with the earlier lines flowering at approximately 71-80 days after planting and the later lines flowering at approximately 80-87 days after planting. Entries were randomized within the block and replicate. These flowering time blocks within replicates were included to account for variation due to flowering time that often contributes to variation in plant height. The inbred lines B73 and PH207 were planted as checks within each block, with five entries of each check per block.

### Manual Plant Height Data Collection

Manual plant height measurements were collected on the same day as drone flights for experimental plots evaluated in this study in 2020 and 2021 (Table S12), and on other experimental plots not included in this study, but in the same field in 2018 and 2019. Manual plant height measurements were collected on 25 randomly distributed plots in 2020 and 15 randomly distributed plots in 2021. Manual plant heights from 144 plots in 2018 and 240 plots in 2019 were collected as previously reported (Tirado *et al*., 2021). Across all four years, within a plot, manual measurements were collected on five representative plants from the center of the plot using a measuring stick. Plant height was measured as the distance between the ground and the uppermost freestanding vegetative part of the plant until reproductive maturity when height was measured to the top of the tassel. These manual measurements were used to determine the relative accuracy of the extracted heights by correlating the manual measurements to the extracted measurements from the same plots (Figure 7, Figure S9).

**Figure 7.**
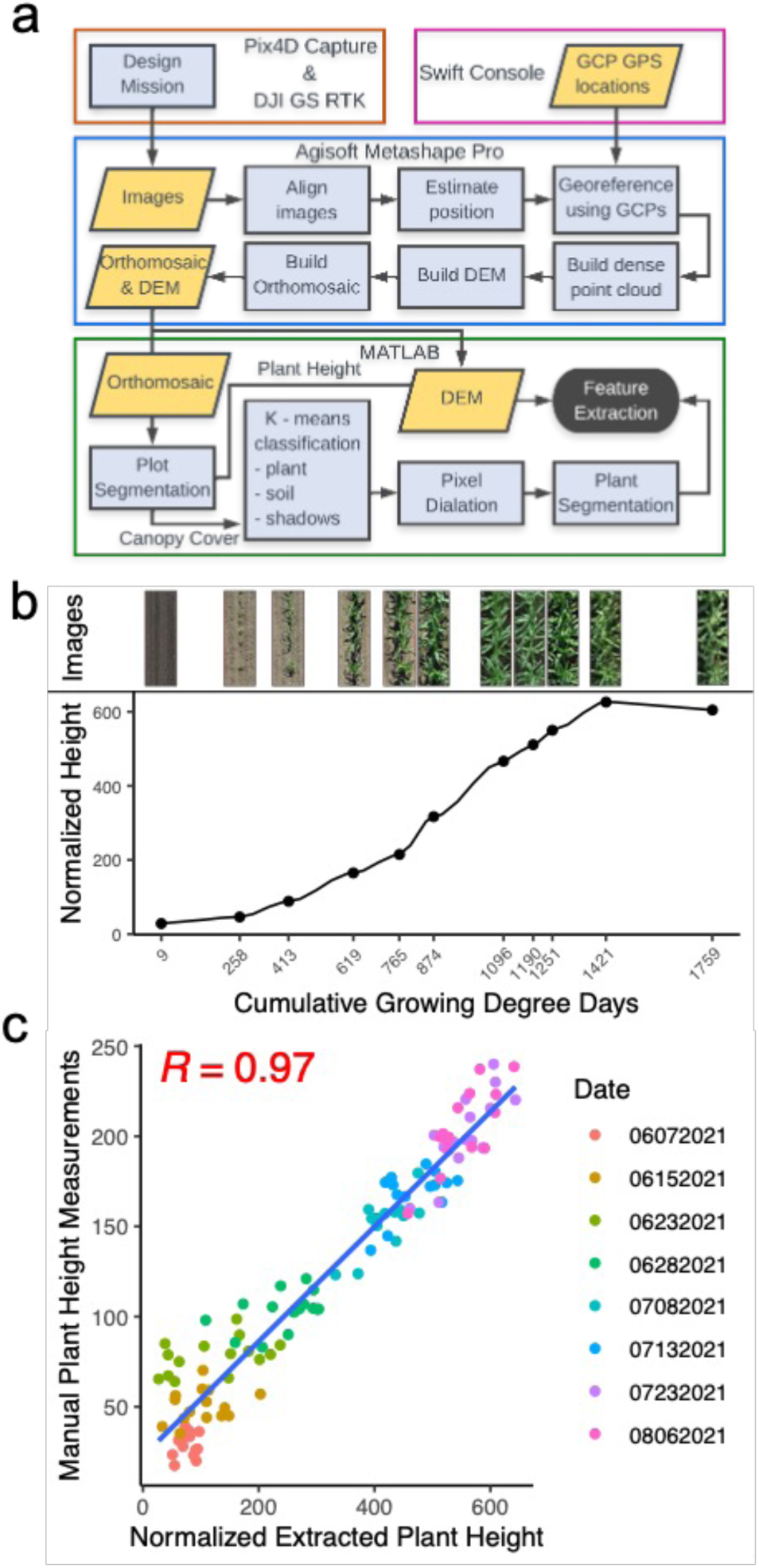
UAV data processing and plant height extraction. (a) Pipeline for image acquisition and feature extraction from UAV images. (b) Extracted plant height normalized across time points for a single example plot with segmented views of the plot at each time point. (c) Pearson correlation between normalized mean extracted plot plant height and mean manual plot plant height measurements across all dates within a year for height extraction validation.

### UAV Data Collection and Processing

The experiment was imaged approximately weekly from planting until plants reached terminal height using a DJI Phantom 4 Advanced drone in 2018 and 2019 and a DJI Phantom 4 RTK drone in 2020 and 2021. Images were collected at an altitude of 30 m above ground to achieve a ground sampling distance of approximately 0.82 cm with 80% front overlap and 80% side overlap to maximize reconstruction efficiency. Flights were collected at 14 timepoints in 2018, 27 timepoints in 2019, 12 timepoints in 2020, and 11 timepoints in 2021 (Table S1). Ground targets of known height and half a meter wide were placed around the border of the field for use as ground control points (GCPs). There were 9 GCPs included in 2018, 12 in 2019, 8 in 2020, and 7 in 2021. The real world coordinates of these GCPs were collected using real time kinematic positioning with a Swift Console (v2.3.17) base station and rover (GNSS compass configuration, n.d.) (Table S13).

### Plant Height Data Extraction From Aerial Images

Images from each flight were processed as previously described (Tirado *et al*., 2020). Briefly, Agisoft Software (Agisoft Metashape Professional v1.7.5) was used to process the images and generate crop surface models and RGB orthomosaics for each flight. QGIS software (QGIS v3.16, 2021) was used for plot boundary extraction by overlaying a grid based on plot size and spacing and exporting plot coordinates. Custom MATLAB scripts for plant height were used to extract height estimates for individual plots using a previously described exposed alley subtraction method (Tirado *et al*., 2020) (Figure 7a, Table S2). All extracted plant heights were normalized across flights by comparing the extracted GCP heights (Figure S1) to the known GCP heights. This was completed by dividing the actual GCP height by the extracted GCP height and then multiplying the value by the extracted plant height (Figure S1a).

The normalized plant heights were subjected to quality control analysis at the field, plot, and individual data point levels. At the field level, visual assessment of heights across the field in heat maps removed four erroneous flight dates in 2019 and an entire second location in 2020 that were not included in the flight counts above. Plots with less than 10 plants were removed from the analysis due to the decrease in extracted plant height accuracy and the competition from neighboring plots affecting the actual plant height. This filtering step removed 325 out of the original 4,160 plots across the four environments. Individual data points (i.e. single plots within a single flight date) were removed if there was a dip in height when the plot height was less than 80% of the previous day and also remained less than the next day, or if the individual data points were classified as a peak when the plot height was more than 120% of the next day while still remaining more than the previous day (n=2,273 data points removed as dips or peaks). Entire plots were removed from a location if the plot height dipped or peaked on 3 or more occasions (n=172 plots removed). In total, 8,238 individual data points (1,201 in 2018, 3,743 in 2019, 2,029 in 2020, and 1,265 in 2021) were removed. After these filtering steps, 3,663 of the original 4,160 plots across the four years had at least 9 individual measurements within the year and were retained for downstream analysis (Table S3).

### Weather Data and Growing Degree Day Unit Calculations

Daily minimum and maximum temperature and precipitation data were collected from the University of Minnesota St. Paul weather station (Station ID: 218450) (Minnesota Department of Natural Resources, n.d.). Growing Degree Day Units (GDDs) were calculated for each date utilizing the equation:*GDD* = [(*Tmax* + *Tmin*)/*2*] − *50* where GDD is the growing degree day units accumulated for a single day and Tmax and Tmin are the maximum and minimum recorded air temperature values in degrees Fahrenheit for the given date. Temperature values above 86°F were adjusted to 86°F and values below 50°F were adjusted to 50°F since corn growth rates do not increase or decrease outside of this range. The cumulative sum of GDDs for data collection dates was calculated as the sum of GDDs for all days between the planting date and the flight date (Figure S1b). Daily data for 18 weather features were obtained for each year using envirotypeR in R v.3.6.2 (R Core Team, 2019) from planting to approximate flowering time at 90 days after planting (Table S5). Correlations between years were calculated by converting the environmental data into pairwise Euclidean distance matrices using the ‘daisy’ function in the ‘cluster’ package v.2.1.4 (Maechler *et al*., 2023) and then using the ‘mantel.rtest’ function in the ‘ade4’ package v.1.7-22 (Dray and Dufour, 2007) in R v.3.6.2 (R Core Team, 2019) to calculate the correlations between each pair of years.

### LOESS Regression Modeling

Separate LOESS regression models were fit to the RGB extracted plant heights for each plot using all flight time points that passed the above filtering criterion using the ‘loess’ function in the ‘stats’ package v.3.6.2 in R v.3.6.2 (R Core Team, 2019). A span of 0.15 to 0.35 was used for regression modeling based on an error matrix developed with 100 folds to determine the best fitting span for each year (Figure S10). Plant height values for every 50 GDDs of the growing season for each model (plot within a year) were predicted using the fitted model and trimmed to 150 to 1450 GDDs. These values were used to calculate the rate of growth between two time points as the slope between the two points (Table S4).

### Analysis of Variance

The variation in extracted plot plant height values and growth rates was partitioned into genotype, year, replicate nested within year, block nested within replicate within year, and genotype by year interaction with all terms treated as fixed effects with the linear model plant height ∼ g + y + y/r + y/r/b + g:y + ε where g is genotype, y is year, r is replicate, b is block, and ε is residual. To test the significance of the sources of variation of the linear model on plant height and growth rate, the ‘Anova’ function from the ‘car’ package v.3.1-1 (Fox and Weisberg, 2019) in R v.3.6.2 (R Core Team, 2019) was used. The ‘tidy’ function in the ‘broom’ package v.1.0.3 (Robinson *et al*., 2024) was used to capture the output of the ANOVA and calculate the percent variance explained by each source of variation in R v.3.6.2 (R Core Team, 2019). Narrow sense heritability (*h*) was calculated using the ‘VarCorr’ function from the ‘nlme’ package v. 3.1-160 (Pinheiro *et al*., 2024) in R v.3.6.2 (R Core Team, 2019).

### Clustering Growth Curves

The LOESS curves were soft clustered within-year using fuzzy c-means clustering using the ‘ppclust’ package v.1.1.0.1 (Cebeci, 2019) in R v.3.6.2 (R Core Team, 2019) (Table S6). The curves were grouped into individual clusters based on the wellness of fit to each cluster, with each curve placed in the group with the highest percentage fit. Comparisons of the clustering between years was completed to determine the consistency of growth curve clustering using an upset plot showing the count of genotypes shared in each cluster type using the ‘upset’ function in the ‘UpSetR’ package v.1.4.0 (Gehlenborg, 2019). A Fréchet distance was calculated between replicates of the same genotype within each year and across years using the ‘TSdist’ package v.3.7.1 (Mori *et al*., 2016) in R v.3.6.2 (R Core Team, 2019) with replicates within a year averaged at each time point to compare across years (Table S8). The genotypes were divided into groups based on the average Fréchet distance and terminal height variance within this dataset. Anything in the lowest 20 percent (n = 100) was considered to be low (Fréchet distance less than 0.330 or plant height variance less than 1150). Anything in the highest 20 percent was considered high (Fréchet distance greater than 0.510 or plant height variance greater than 4075).

### Genome-Wide Association Studies

Genome-wide association studies were completed as previously described (Renk *et al*., 2021; Burns *et al*., 2021). In brief, GAPIT v.3 (Wang and Zhang, 2021) was used to transform genomic data into numeric format, generate a genotypic map dataset, and a PCA covariates dataset. SNP data was obtained from a previous study (Qiu *et al*., 2021) and filtered as previously described for minor allele frequency, missing data, and LD within 10 kb (Renk *et al*., 2021). A separate GWAS was performed for each of the predicted GDD time points, derived growth rates, and normalized extracted terminal heights in each year using BLUPs extracted from the fixed effects model described above within each year. The genetic map, filtered numeric genomic dataset, and filtered BLUP datasets were used to permute a suggested p-value for each GWAS using the ‘FarmCPU.P.Threshold’ function in FarmCPU v.1.02, (Liu *et al*., 2016) in R v.4.0.4 (R Core Team, 2019) as previously described (Renk *et al*., 2021). SNPs in linkage disequilibrium with each other were identified using PLINK v.1.90b6.10 (Purcell *et al*., 2007; Chang *et al*., 2015; Shaun Purcell, n.d.). SNP effect over time was plotted in a heatmap using ‘heatmap.2’ from the ‘gplots’ package v.3.1.3 (Warnes *et al*., 2024) (R Core Team, 2019) and the SNP effects were clustered using the default ‘hclust’ function from heatmap.2 (Table S9). The nearest gene to a SNP was identified relative to the Zm-B73-REFERENCE-GRAMENE-4.0 Zm000014 Gene Set from MaizeGDB (Maize B73 RefGen_v4) (Monaco *et al*., 2013). Functional annotations for this gene set were downloaded from Gramene (ftp://ftp.gramene.org/pub/gramene/CURRENT_RELEASE/gff3/zea_mays/gene_function/B73v 4.gene_function.txt) on 8-Nov-2018. Overlap with significant regions from previous studies (Mazaheri *et al*., 2019; Zhang *et al*., 2019; Adak, Murray, Anderson, *et al*., 2021; Wang *et al*., 2019) was completed using the ‘window’ function in bedtools version 2.31.1 (Quinlan and Hall, 2010) with SNPs within 100 kb considered overlapping.

### Canopy Cover Data Analysis

The same flights that were processed for plant height in this study were previously processed for canopy cover using the same crop surface models, RGB orthomosaics, and plot boundaries described above (Cooper *et al*., 2024). LOESS regression models were fit using the same methods described above for plant height and point and growth rate values in 50 GDD windows were predicted from the models (Table S11).

As with plant height, a Fréchet distance was calculated between replicates of the same genotype within years and between the same genotype across years for canopy cover using the ‘TSdist’ package v.3.7.1 (Mori *et al*., 2016) in R v.3.6.2 (R Core Team, 2019) (Table S8). Soft clustering was completed within-year using fuzzy c-means clustering using the ‘ppclust’ package v.1.1.0.1 (Cebeci, 2019) in R v.3.6.2 (R Core Team, 2019) (Table S6). Similarities in clustering were evaluated in an upset plot using the ‘UpSetR’ package v1.4.0 (Gehlenborg, 2019) in R v.3.6.2 (R Core Team, 2019). Z-scores were calculated for both plant height and canopy cover using the following equation: *z* = (*x* − *μ*)/sd where z is the z-score for the individual plot, x is the individual plot plant height or canopy cover value, μ is the average value across all retained data points for plant height or canopy cover respectively, and sd is the standard deviation across all retained data points for plant height or canopy cover respectively in order to make plant height and canopy cover comparable in scale to one another. Pearson correlations and Fréchet distances between plant height and canopy cover in the same experimental plot were completed using the ‘cor’ function in the ‘stats’ package v.3.6.2 and the ‘TSdist’ package v.3.7.1 (Mori *et al*., 2016) in R.v.3.6.2 (R Core Team, 2019).

### Data and Code Availability

Scripts and files used to generate and analyze data are available on GitHub at https://github.com/HirschLabUMN/WiDiv_Drone_Height.git. All data including the UAV-derived plot height values, UAV-derived canopy cover values, orthomosaics, DEMs, plot boundary files, and mask files for each date of UAV data collection, cumulative GDDs calculated for each date of data collection, manual height data, and weather data has been made available at the Digital Repository for U of M (DRUM) at https://doi.org/10.13020/SKJN-QX31.

## Acknowledgments

We thank the Minnesota Supercomputing Institute at the University of Minnesota (http://www.msi.umn.edu) for providing resources that contributed to the research results reported in this article. This work was supported in part by the Minnesota Corn Research and Promotion Council and NSF IOS-1546727. D.D.S was funded by the University of Minnesota MnDRIVE Global Food Ventures Graduate Fellowship. S.B.T. was funded by the University of Minnesota Graduate Opportunity Fellowship, the University of Minnesota APS Metric Funds Fellowship, and the Bayer/University of Minnesota Multifunctional Agriculture Initiative Graduate Student Fellowship. J.C. was funded by the Bayer/University of Minnesota Multifunctional Agriculture Initiative Graduate Student Fellowship.

