## Supplemental Figures 1-10 for "Temporally resolved growth patterns reveal novel information about the polygenic nature of complex quantitative traits"

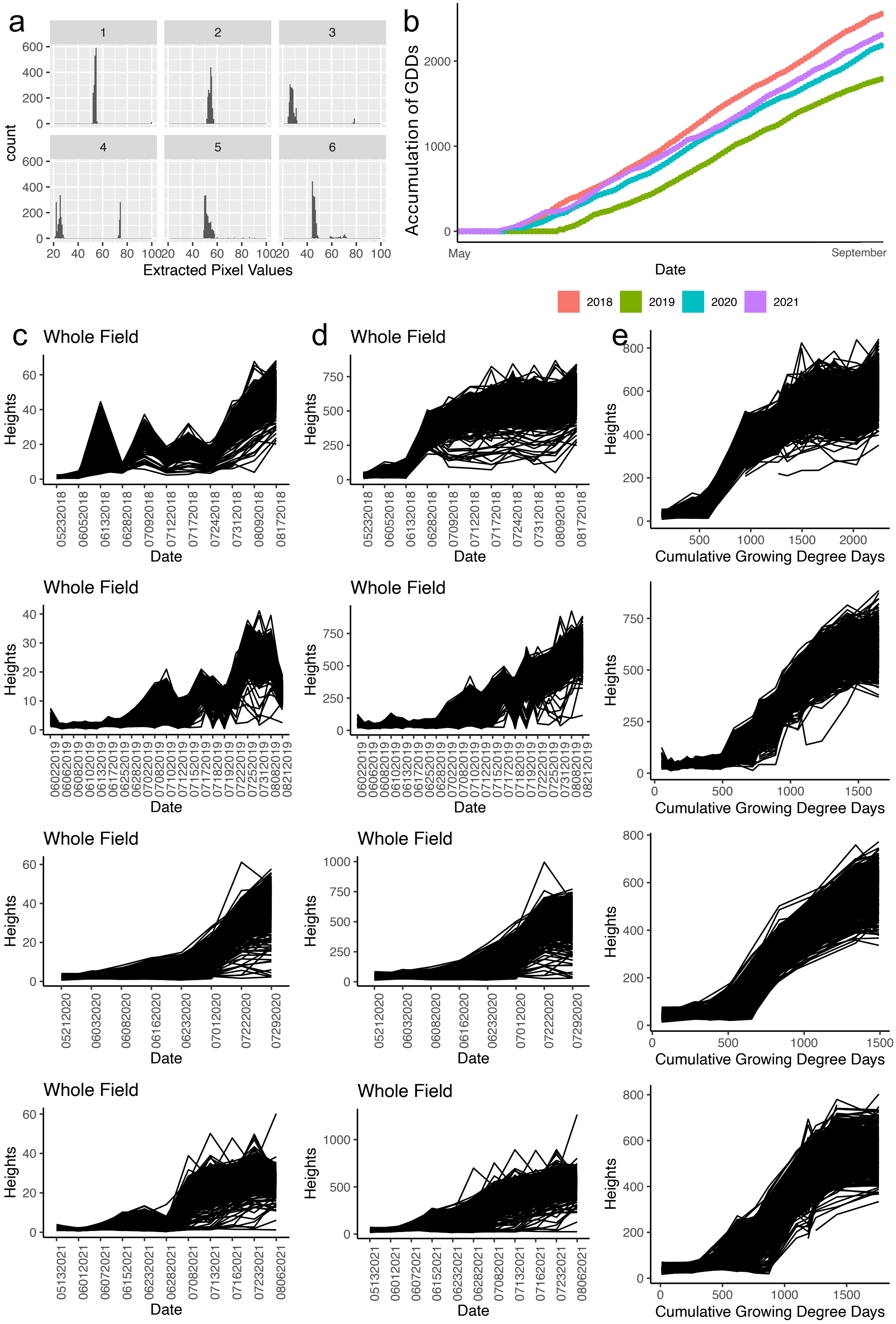


**Figure S1. Normalization of extracted plant height values.** (a) Example values extracted from GCP bounding boxes to identify height values for the ground and tops of the GCPs (06022019). (b) Accumulation of growing degree days (GDDs) throughout the growing season for each year. Extracted plant height values across the growing season for all plots with (c) raw data (before any cleaning), (d) normalized data (before cleaning but normalized across dates using GCP heights), and (e) clean data (normalized data with erroneous plots removed before analysis). Dates on the x-axis are in the form MMDDYYYY.

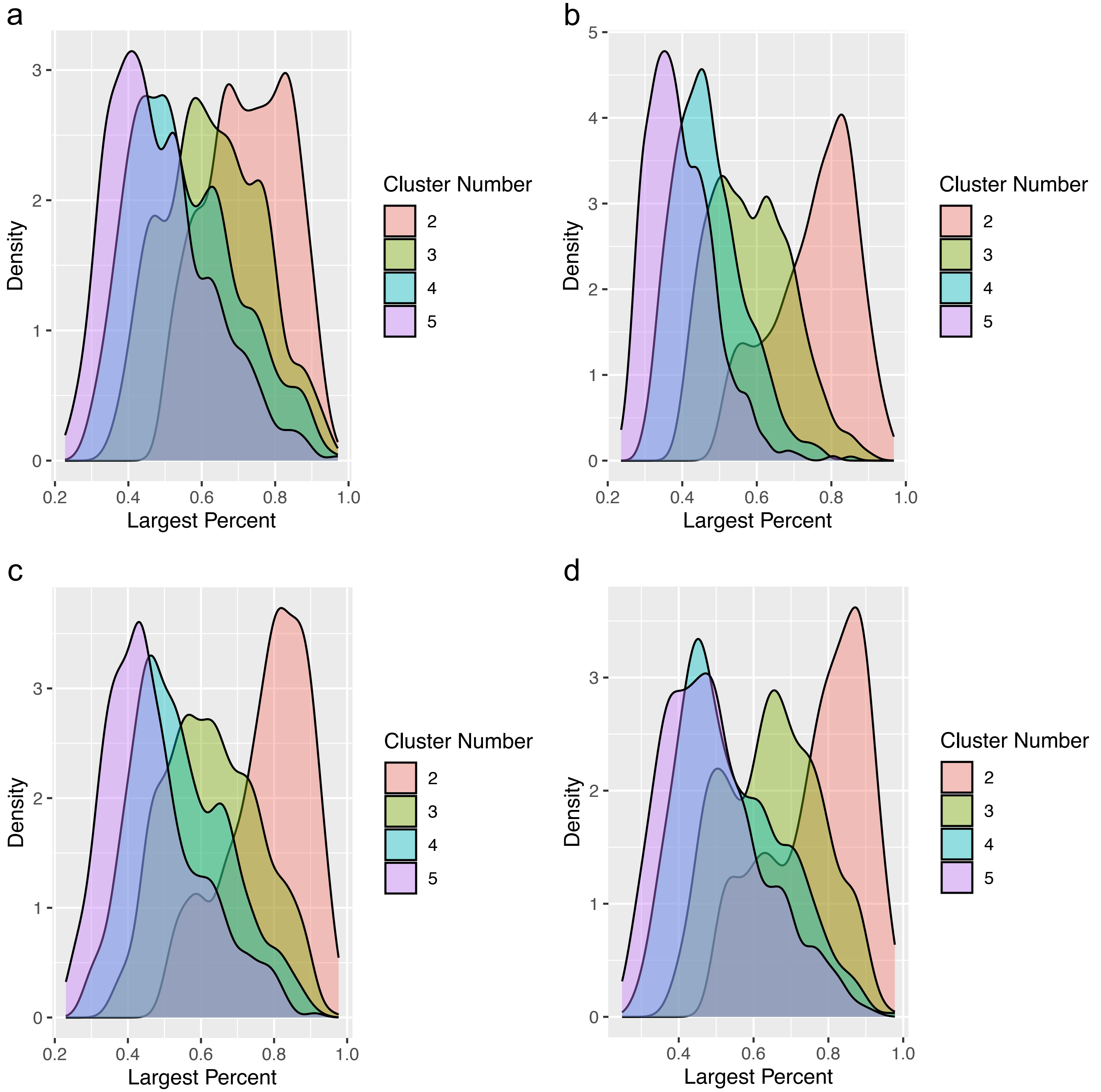


**Figure S2. Density of largest percentages for determination of number of clusters to use for Fuzzy c-means clustering.** The density of the largest value of wellness of fit for each curve to any single cluster when the curves are broken into 2, 3, 4, or 5 clusters with the value of the largest percentage of fit on the x-axis. These graphs were used to determine how many clusters fit the data best. (a) 2018 (b) 2019 (c) 2020 (d) 2021.

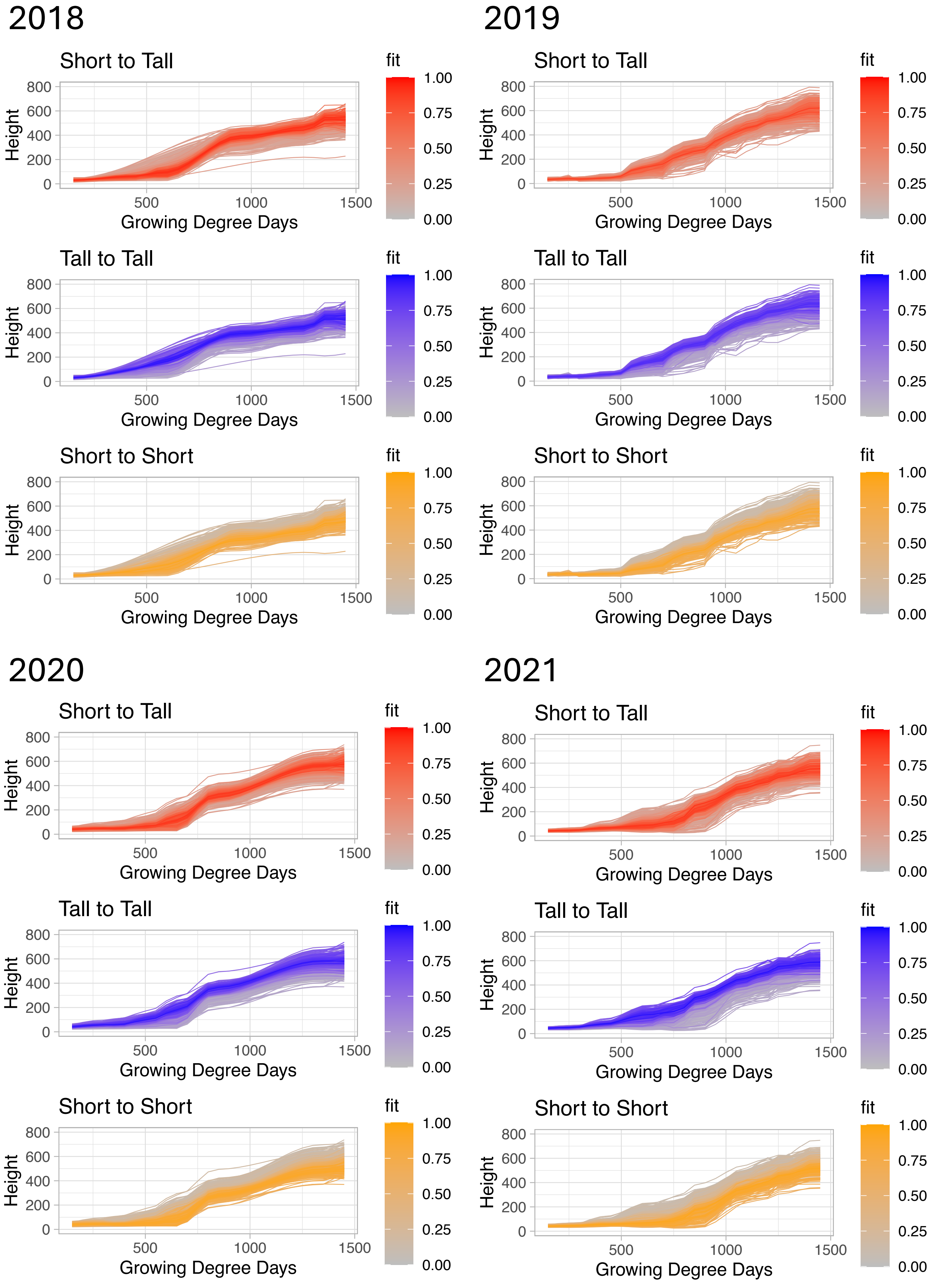


**Figure S3. Fuzzy c-means clustering of plant height growth curve values.** Fuzzy c-means clusters of LOESS growth curves with shading equating wellness of fit for each curve into the specified cluster separately for each year.

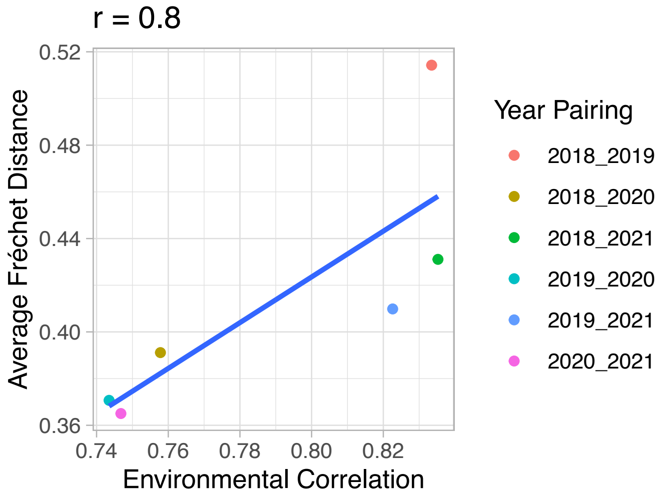


**Figure S4. Pearson correlation between environmental similarity and average Fréchet distance.** Fréchet distance between years for all genotypes with data in both years were averaged within each pair of years. Environmental correlations were calculated based on daily values for 18 weather parameters from planting to approximate flowering at 90 days after planting.

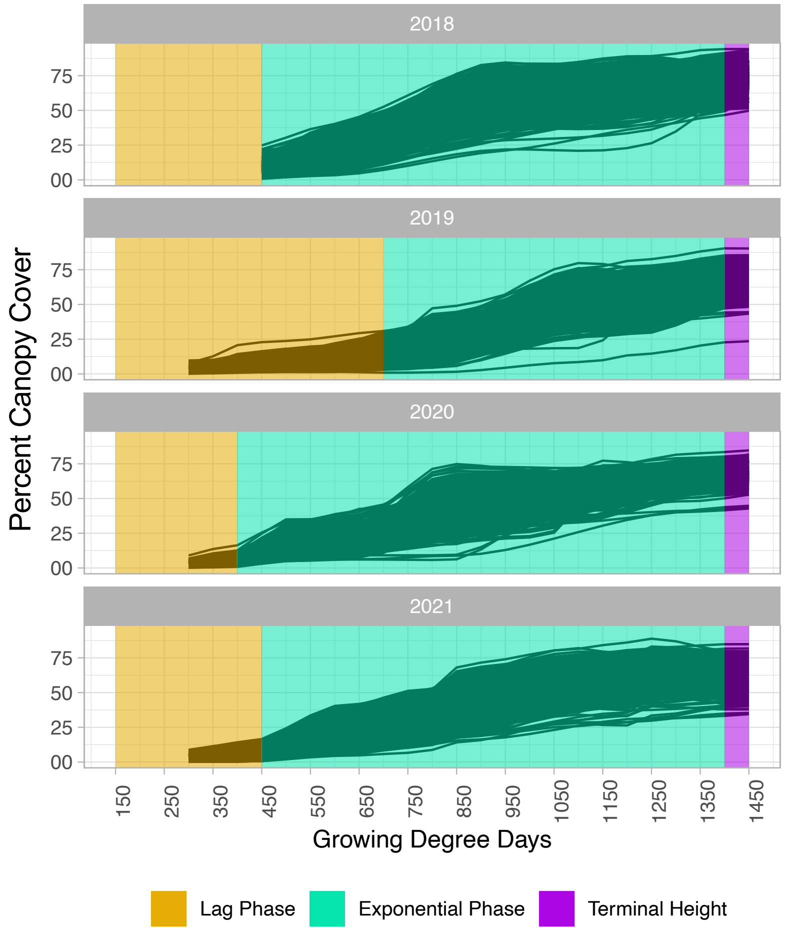


**Figure S5. Temporal canopy cover growth curves.** LOESS curves of percent canopy cover broken into three phases of the growing season based on the performance of genotypes in each year.

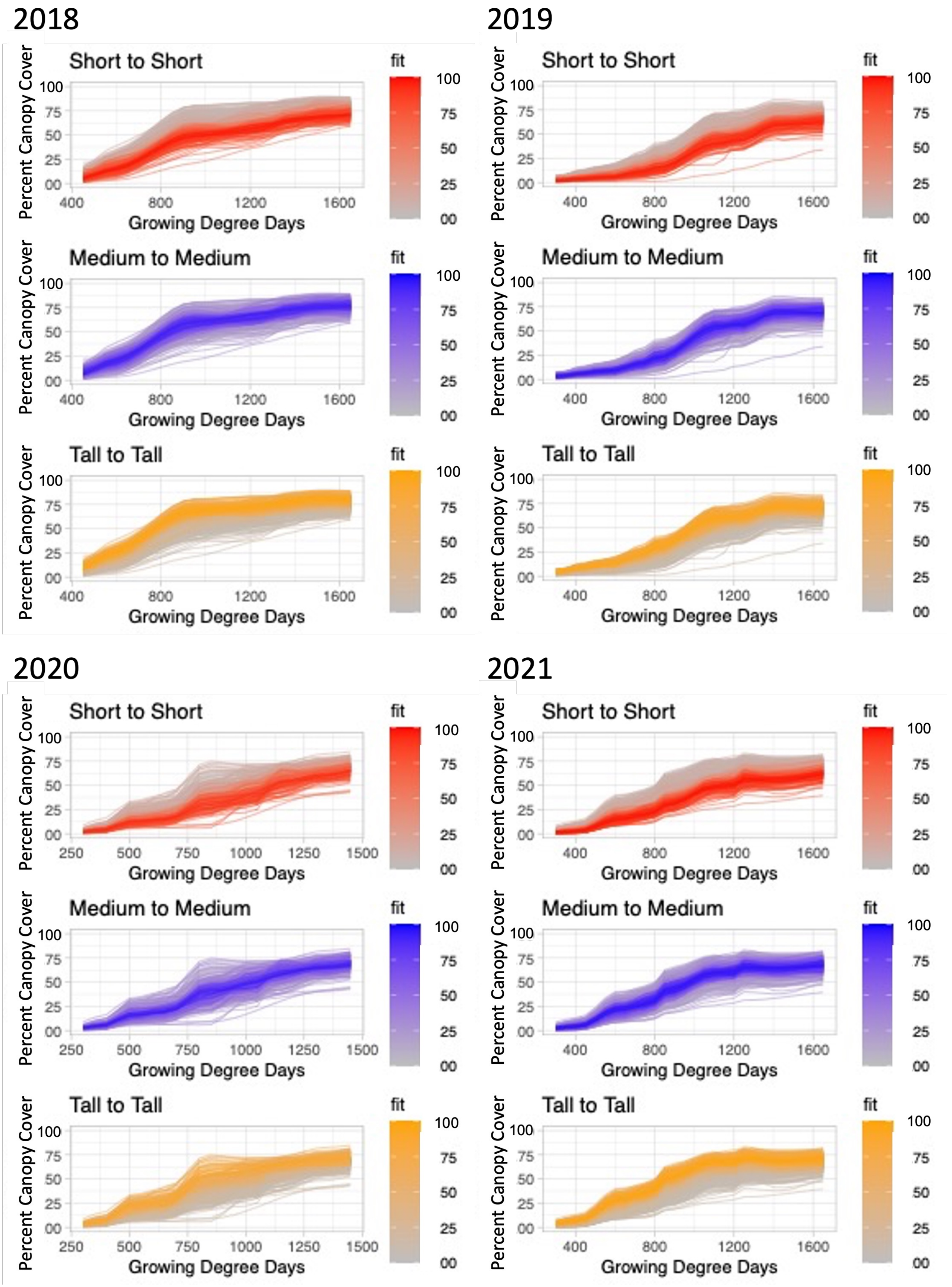


**Figure S6. Fuzzy c-means clustering of canopy cover growth curve values.** Fuzzy c-means clusters of LOESS growth curves with shading equating wellness of fit for each curve into the specified cluster separately for each year.

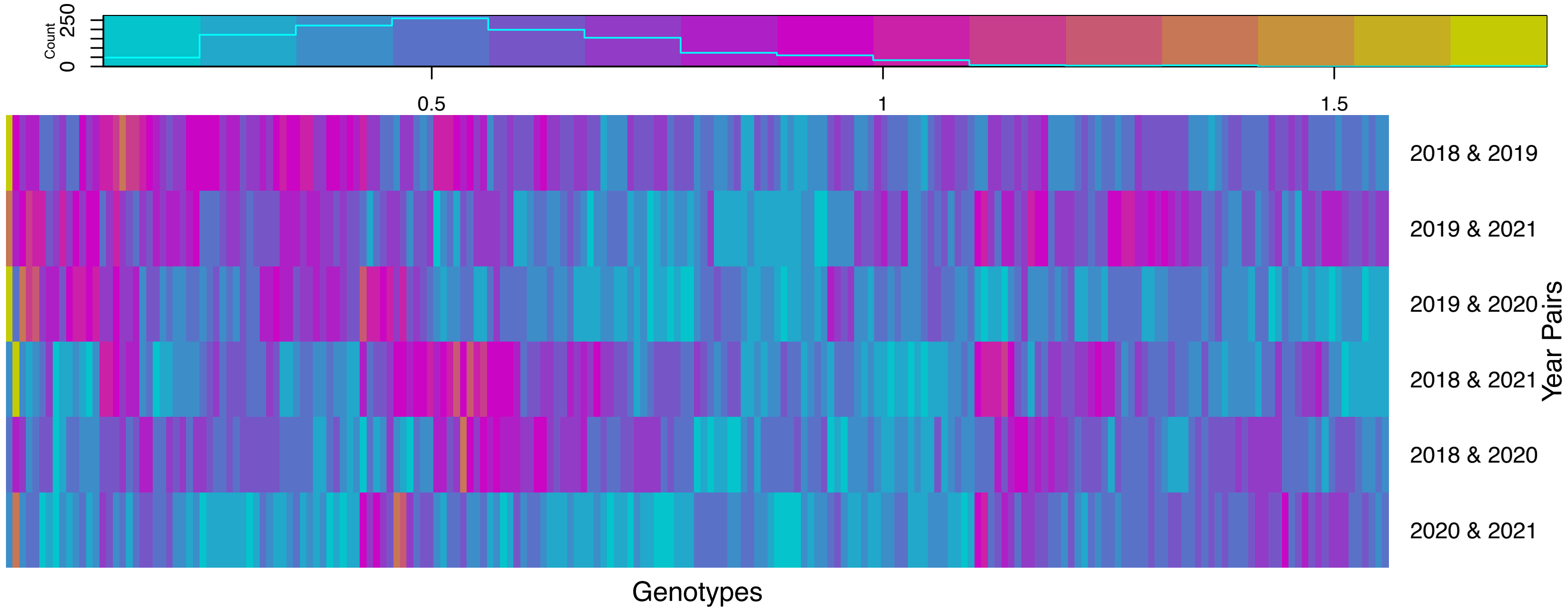


**Figure S7. Fréchet distances for each genotype present in all four years.** Pairwise Fréchet distance values comparing canopy cover growth curves of the same genotype across different years.

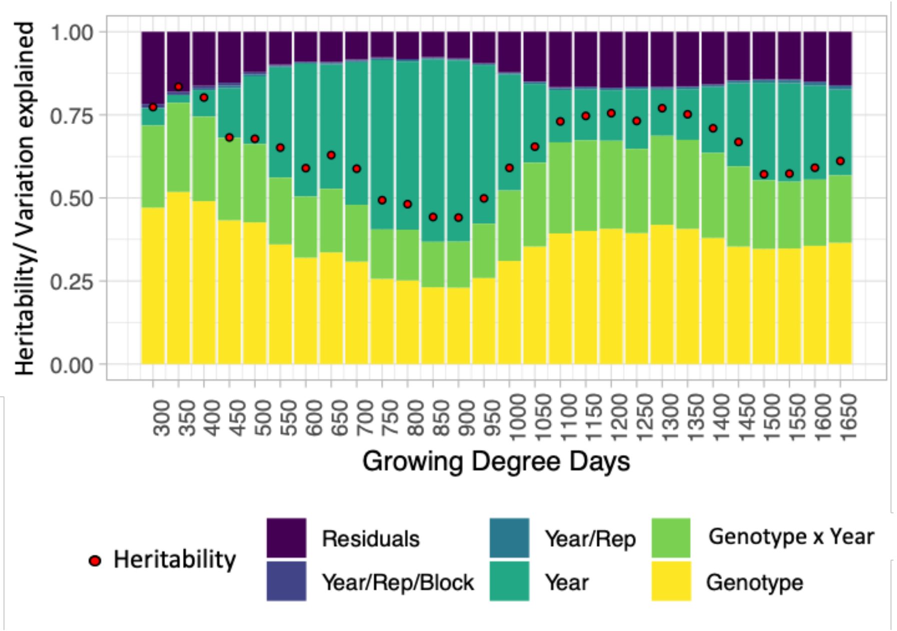


**Figure S8. Temporal canopy cover analysis of variance.** Percent variance explained and heritability at each canopy cover time point throughout the growing season.

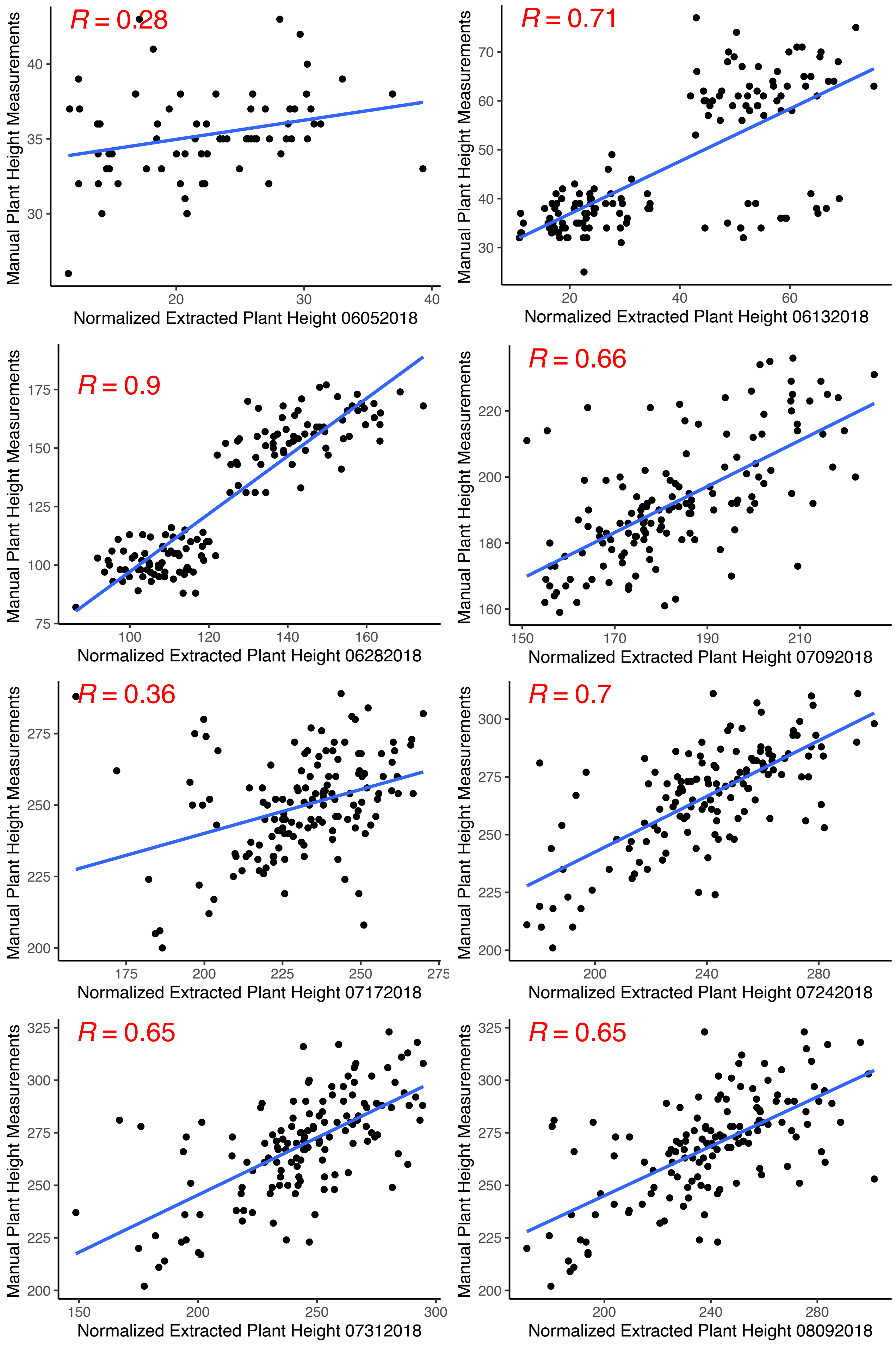

R = 0.28
Manual Plant Height Measurements
Normalized Extracted Plant Height 06052018
R = 0.71
Manual Plant Height Measurements
Normalized Extracted Plant Height 06132018
R = 0.9
Manual Plant Height Measurements
Normalized Extracted Plant Height 06282018
R = 0.66
Manual Plant Height Measurements
Normalized Extracted Plant Height 07092018
R = 0.36
Manual Plant Height Measurements
Normalized Extracted Plant Height 07172018
R = 0.7
Manual Plant Height Measurements
Normalized Extracted Plant Height 07242018
R = 0.65
Manual Plant Height Measurements
Normalized Extracted Plant Height 07312018
R = 0.65
Manual Plant Height Measurements
Normalized Extracted Plant Height 08092018

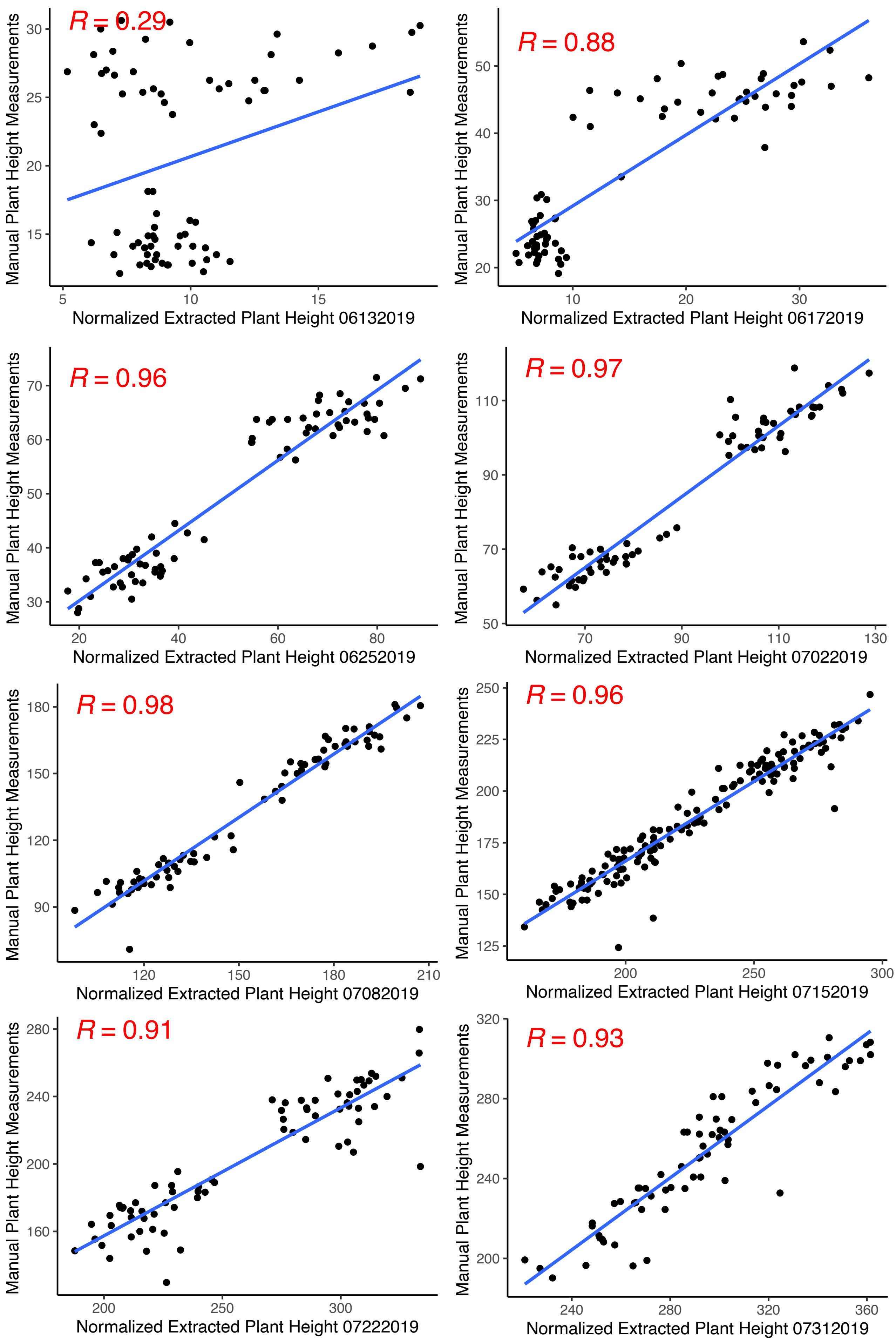

R = 0.29
Manual Plant Height Measurements
Normalized Extracted Plant Height 06132019
R = 0.88
Manual Plant Height Measurements
Normalized Extracted Plant Height 06172019
R = 0.96
Manual Plant Height Measurements
Normalized Extracted Plant Height 06252019
R = 0.97
Manual Plant Height Measurements
Normalized Extracted Plant Height 07022019
R = 0.98
Manual Plant Height Measurements
Normalized Extracted Plant Height 07082019
R = 0.96
Manual Plant Height Measurements
Normalized Extracted Plant Height 07152019
R = 0.91
Manual Plant Height Measurements
Normalized Extracted Plant Height 07222019
R = 0.93
Manual Plant Height Measurements
Normalized Extracted Plant Height 07312019

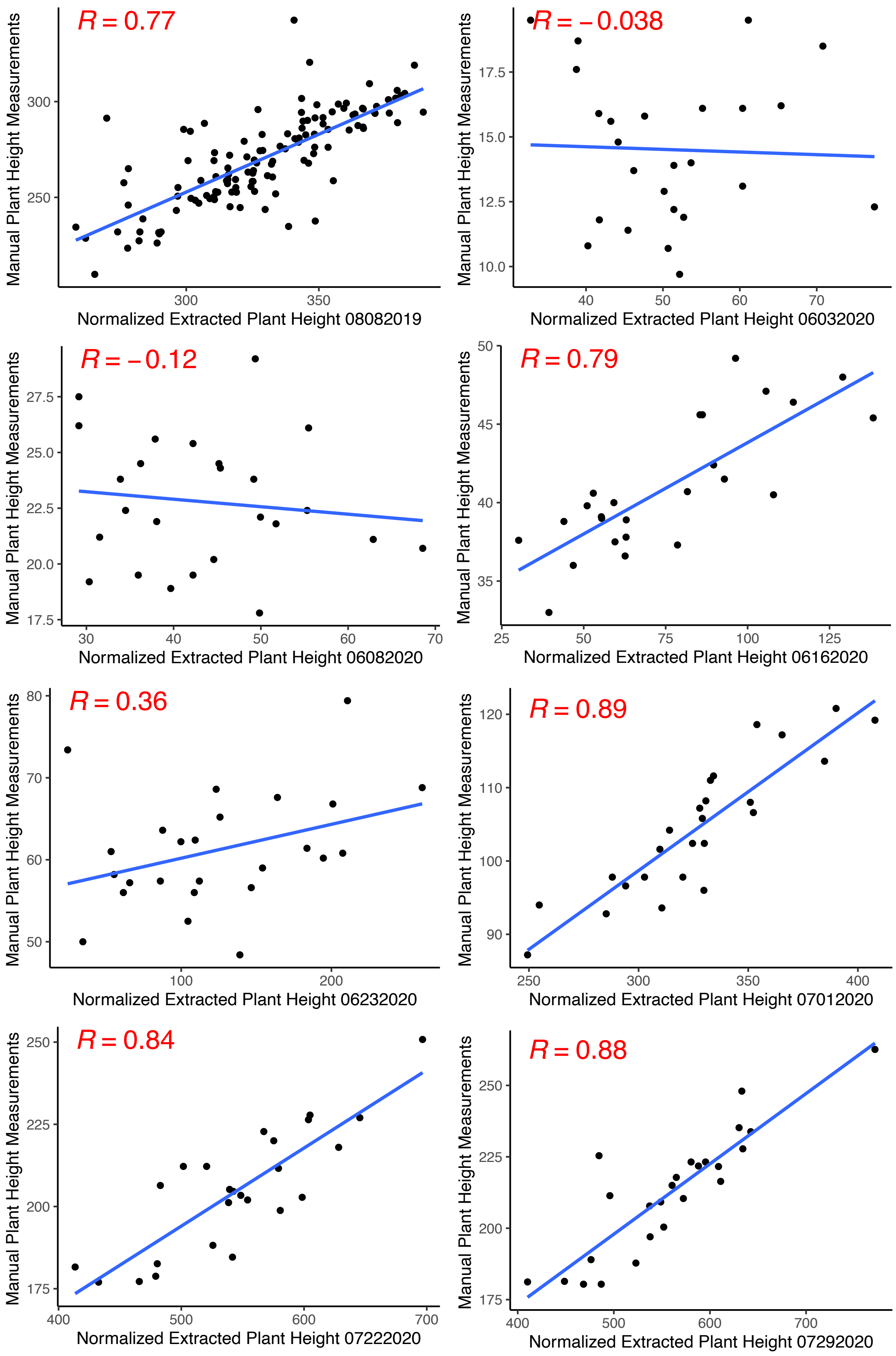
R = 0.77
Manual Plant Height Measurements
250
300
300
350
Normalized Extracted Plant Height 08082019
R = −0.038
Manual Plant Height Measurements
10.0
12.5
15.0
17.5
40
50
60
70
Normalized Extracted Plant Height 06032020
R = −0.12
Manual Plant Height Measurements
17.5
20.0
22.5
25.0
27.5
30
40
50
60
70
Normalized Extracted Plant Height 06082020
R = 0.79
Manual Plant Height Measurements
35
40
45
50
25
50
75
100
125
Normalized Extracted Plant Height 06162020
R = 0.36
Manual Plant Height Measurements
50
60
70
80
100
200
Normalized Extracted Plant Height 06232020
R = 0.89
Manual Plant Height Measurements
90
100
110
120
250
300
350
400
Normalized Extracted Plant Height 07012020
R = 0.84
Manual Plant Height Measurements
175
200
225
250
400
500
600
700
Normalized Extracted Plant Height 07222020
R = 0.88
Manual Plant Height Measurements
175
200
225
250
400
500
600
700
Normalized Extracted Plant Height 07292020

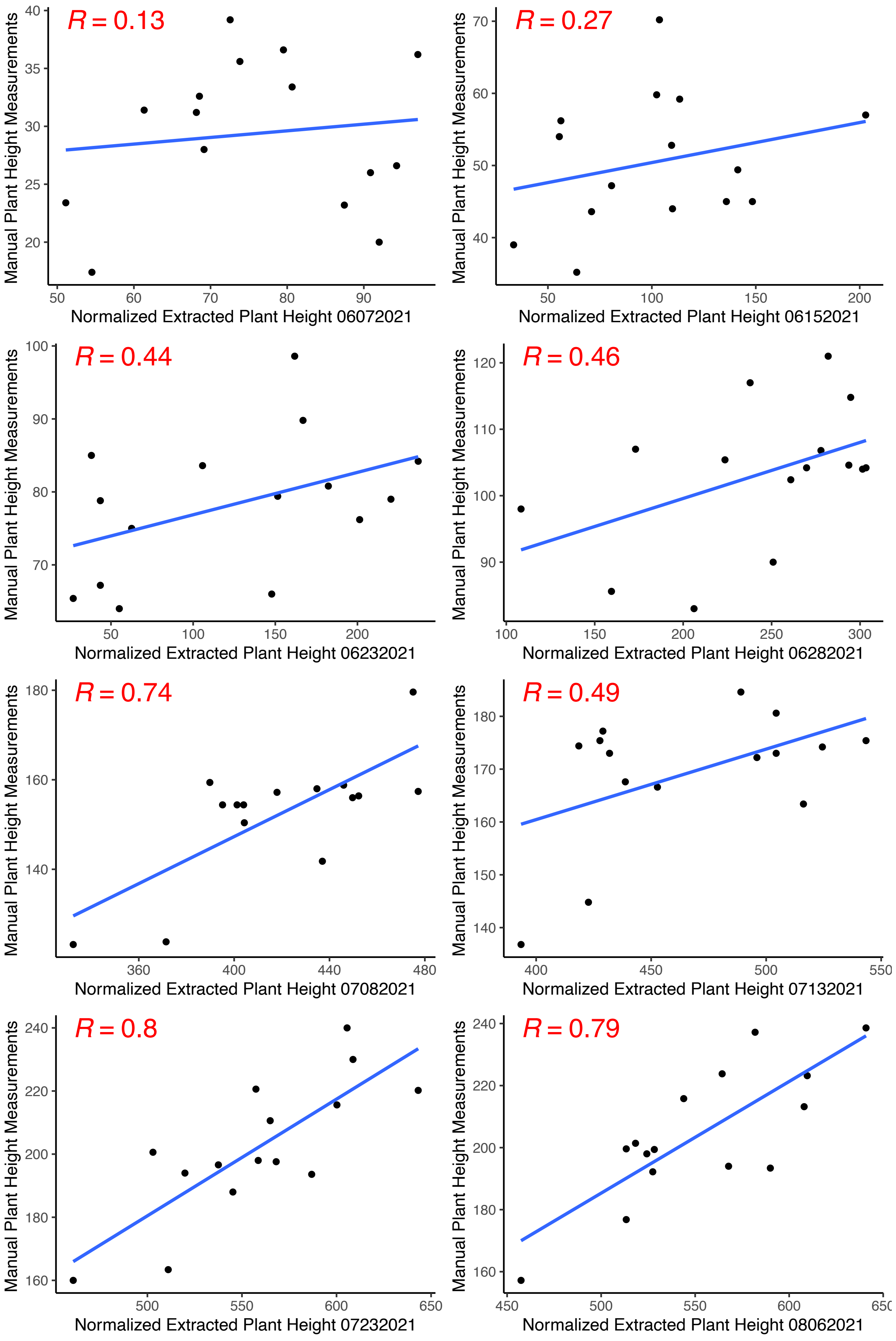


**Figure S9. Validation of extracted plant height.** Pearson correlation of normalized mean extracted plot plant height to mean manual plot plant height measurements. The date of each flight is indicated in the x-axis label in the form MMDDYYYY.

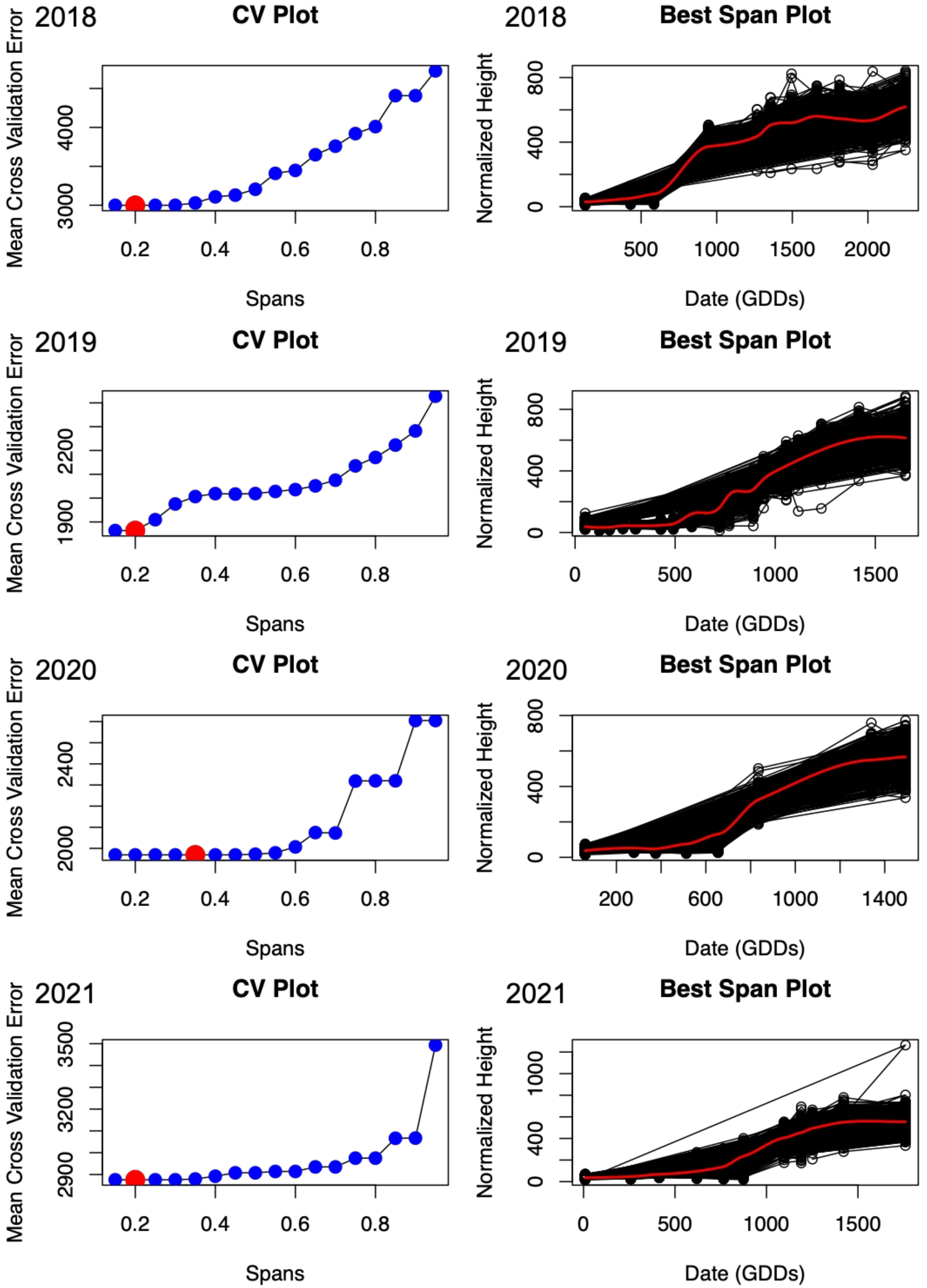


**Figure S10. Span choices for LOESS curve fitting each year.** Mean cross validation error for each possible span from 0.15 to 0.95 for each year with the best span for each year red (left plots). Growth rates with loess curves fit using the best span (right plots).
